# Size-Dependent Mechanoadaptation Enables Migration with Oversized Parasitic Cargo

**DOI:** 10.1101/2024.05.03.592378

**Authors:** Mauricio J.A. Ruiz-Fernandez, Jianfei Jiang, Alice Battistella, Armina Mortazavi, Sterre van Wierst, Bingzhi Wang, Artur Kuznetcov, Rüya Aslan, Jack Merrin, Daan Vorselen, Jochen Guck, Markus Meissner, Javier Periz, Benedikt Sabass, Jörg Renkawitz

## Abstract

Cell migration is fundamental to development, homeostasis, and immunity, and requires cells to traverse space-limited microenvironments by deforming intracellular organelles such as the nucleus. Although nuclear mechanics are established as a rate-limiting factor during confined migration, how cells transport bulky foreign cargo remains unclear. Here, using *Toxoplasma gondii*, a disease-causing parasite that hijacks motile immune cells for dissemination while replicating into large intracellular assemblies, we discover that parasitic cargoes exceed the size and stiffness of host nuclei, and therefore constitute a primary bottleneck for migration. We uncover a mechanoadaptive response to parasite size, characterized by size-dependent intracellular repositioning, myosin-II contractility, and bleb-based protrusions. This reprogramming of forces and subcellular architecture enables the transport of oversized, mechanically rigid cargo through space-limited microenvironments. These findings establish a mechanobiological principle of how cells size-dependently adapt forces and their subcellular architecture to accommodate intracellular objects that challenge the physical limits of cell migration.

## Introduction

Cell movement is essential in biology, impacting processes such as organismic development, tissue maintenance, and immune responses. Within multicellular organisms, motile cells face the challenge of navigating complex local microenvironments with spaces substantially smaller than the cells themselves^1^. As a result, motile cells must continuously deform their cell body, as well as their intracellular organelles, such as the nucleus^2,3^. Since the nucleus is typically the stiffest and largest organelle within a cell, its deformation during movement through space-limiting microenvironments is rate-limiting for transmigration^4,5^. To overcome this challenge, motile cells rely on forces generated by their cytoskeleton, which propel themselves and their organelle cargo through these constraining environments^6–8^.

Immune cells face the challenge of constantly transmigrating through space-limiting microenvironments, as they move along long migratory tracks during their surveillance and effector responses^9,10^. To alleviate the physical burden of their nuclear cargo along these trafficking routes, immune cells frequently possess nuclei with lower stiffness than non-immune cells^11^. Still, their nucleus is stiffer when compared to other subcellular organelles, and its deformation represents a major bottleneck^5^. To enable their rapid movement, immune cells employ dedicated cytoskeleton networks that accurately position their nucleus intracellularly^12^ and squeeze it through narrow pores^6,9^. However, it is not understood how immune cells transport foreign cargoes along their challenging trafficking routes, such as phagocytosed cell corpses, microplastics, and pathogens^13–15^, particularly when these cargoes exceed the size of their own organelles.

One example of such a cargo is the human pathogen *Toxoplasma gondii*, an obligate intracellular parasite that infects a large part of the human population and causes toxoplasmosis, which can lead to encephalitis in immunocompromised patients and fetal abnormalities during pregnancy^16^. Upon uptake, the parasite efficiently infects immune cells in the host gut and arrives in lymph nodes within a few days, which represent the homing destination of many immune cells, as well as in distant organs like the heart and brain^17,18^. This fast dissemination of the parasite relies on two complementary behaviors: (1) the extracellular motility of the parasite itself, and (2) the hitchhiking by motile host cells for its transport as an intracellular cargo. The extracellular motility of the parasite is essential for invading into and egressing from host cells, leading to local dissemination^19,20^. In addition, hitchhiking motile host cells is a second, complementary mechanism for long-range parasitic dissemination. Notably, *T. gondii* efficiently infects host immune cells, thereby hijacking their ability to rapidly move between distant tissue locations^15,21^. Moreover, the parasite induces a highly motile mode of immune cells upon infection^22–26^, which further facilitates parasite dissemination^27,28^. Thus, the parasite not only hides within immune cells to avoid immune detection, but also turns these cells into its own transport shuttles^15^. However, parasite replication within the parasitophorous vacuole^29^ generates large intracellular parasitic objects. This raises the unresolved paradox of how rapidly migrating immune cells efficiently traverse constraining microenvironments while carrying foreign, bulky cargoes.

Here, we discover that the parasitic cargo represents a novel and primary bottleneck for cell migration of infected immune cells, whose transport is even more challenging than the bulky host nucleus. Using custom-made bioengineering, advanced imaging, mechanical force measurements, computer simulations, and genetic engineering of parasites and immune cells, we establish the underlying mechanical and cellular principles that enable efficient immune cell migration despite the physical challenges posed by the foreign cargo. Our findings thereby reveal a new size-dependent mechano-adaptation strategy that redistributes forces and the subcellular architecture.

## Results

### Efficient Transport of Large Assemblies of Intracellular Parasites Through Dense Environments

To address the paradox of how motile immune cells transport large intracellular parasitic cargoes through constrained environments, we employed *Toxoplasma gondii* as a well-established eukaryotic parasite model that grows intracellularly^19,22,23,30,31^, and mouse dendritic cells (DCs) as a well-established model for motile immune cells that are infected by *Toxoplasma* and move with an amoeboid migration strategy^32–34^. Upon timed infection, we loaded parasite-infected DCs into straight tunnel-like microchannels with a defined width of 8 micrometers and height of 5 micrometers (**Figure 1A**), providing a constrained microenvironment that mimics confining tissues and narrow capillaries^35–38^. Using simultaneous imaging of hundreds of microchannels^39^ allowed us to precisely quantify the migratory velocity of parasite-infected DCs within these constraining microenvironments. Importantly, by using genetically engineered *Toxoplasma* parasites that stably encode a Halo-tagged version of the major surface antigen SAG1 for fluorescent visualition^40,41^ (**Figure 1A**), we were able to track the velocity of the infected host cell while mapping the parasitic replication stage and thus the intracellular size of the parasitic cargo (**Figures 1B** and **C**).

**Figure 1.**
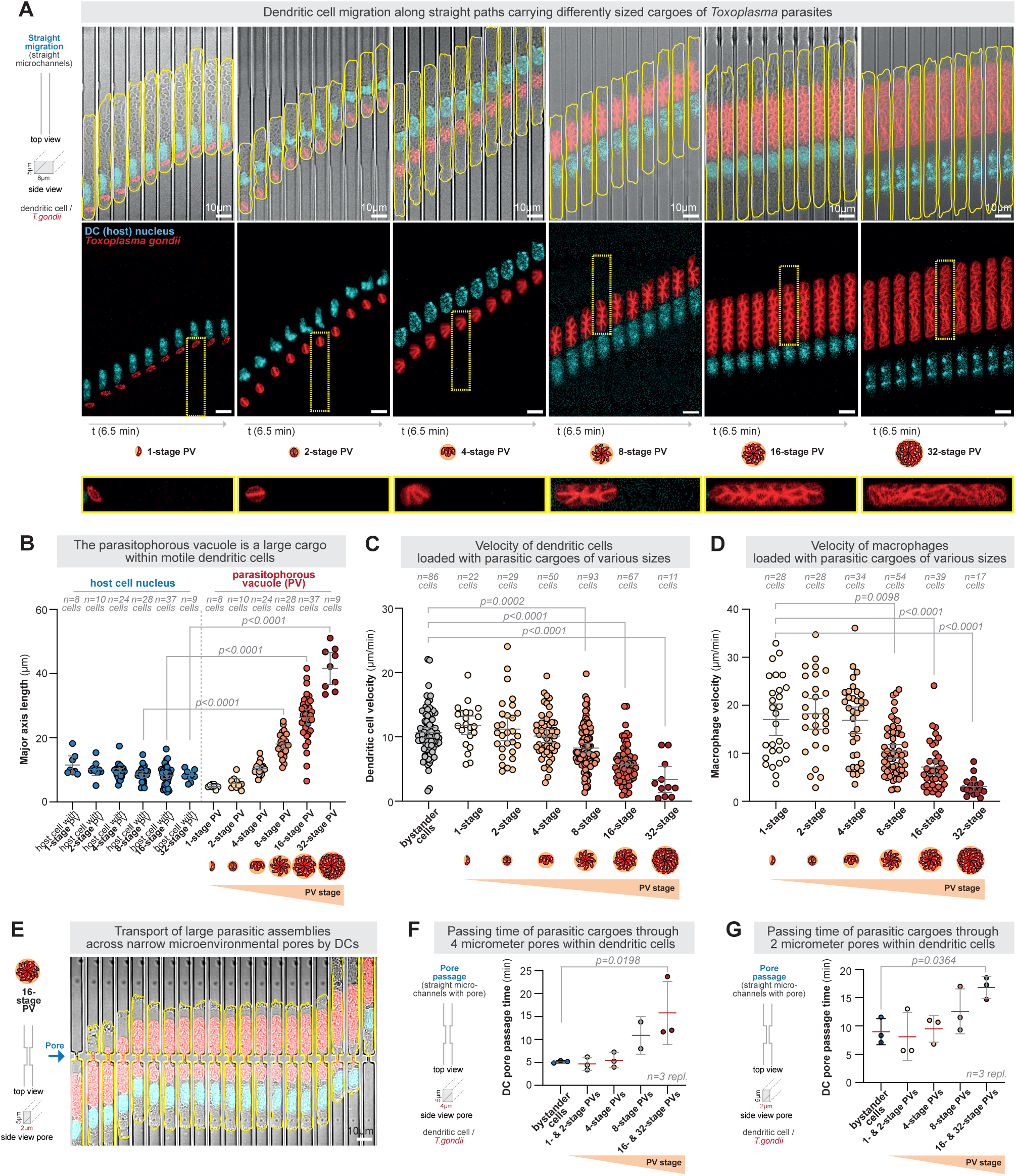
Efficient transport of large assemblies of intracellular parasites by immune cells through dense environments. **(A)** Representative *Toxoplasma gondii*-infected dendritic cells (DCs) migrating in linear microchannels are shown as kymographs over 6.5 minutes. *Toxoplasma gondii* is shown in red, the host cell shape in yellow, and the DC nucleus in cyan (Hoechst). Note the different replication stages (1- to 32-stage) and respective sizes of the parasitic cargo within the motile host cell (see also the zoom-ins in the yellow boxes). **(B)** Quantification of the length of the parasitophorous vacuole in comparison to the host cell nucleus during DC migration along linear microchannels. Note that a 4-stage PV equals the length of the host nucleus, while 8-, 16, and 32-staged PVs exceed the size of the host nucleus. N = 8 cells (1-stage PV), 10 cells (2-stage PV), 24 cells (4-stage PV), 28 cells (8-stage PV), 37 cells (16-stage PV), and 9 cells (32-stage PV) from 3 independent biological replicates. Unpaired t-tests. Data are the mean±95% CI. **(C)** DC velocity in relation to the replication stage of the parasite cargo. N = 86 cells (non-infected bystander DCs), 22 cells (1-stage PV), 29 cells (2-stage PV), 50 cells (4-stage PV), 93 cells (8-stage PV), 67 cells (16-stage PV), and 11 cells (32-stage PV) from 3 independent biological replicates. Kruskal-Wallis multiple comparison test on bystander cells. Data are the mean±95% CI. **(D)** Macrophage velocity in relation to the replication stage of the parasite cargo. N = 28 cells (1-stage PV), 28 cells (2-stage PV), 34 cells (4-stage PV), 54 cells (8-stage PV), 39 cells (16-stage PV), and 17 cells (32-stage PV) from 4 independent biological replicates. Kruskal-Wallis multiple comparison test on 1-stage infected cells. Data are the mean±95% CI. See Figure S1A for representative image examples, Figure S1B for parasitic sizes within macrophages, and Figure S1D for passing times through 2 micron pores within macrophages. **(E)** Representative dendritic cell carrying a 16-stage *Toxoplasma* cargo across a 2-micron-wide pore within a microchannel. *Toxoplasma gondii* is shown in red, the host cell shape in yellow, and the DC nucleus in cyan (Hoechst). **(F)** Passing times of parasitic cargoes of different sizes during DC migration through 4 micrometer-sized pores. See Figure S1C for representative images. One-way ANOVA with multiple comparisons on bystander cells. Data are the mean±SD, N = 3 biological replicates. **(G)** Passing times of parasitic cargoes of different sizes during DC migration through 2 micrometer-sized pores, as shown in Figure 1E. One-way ANOVA with multiple comparisons on bystander cells. Data are mean±SD, N = 3 biological replicates.

Dendritic cells infected with small 1-stage parasites rather showed faster migration velocities than uninfected bystander cells (**Figure 1C**), consistent with previous findings that identified a high mobility of *Toxoplasma*-infected DCs^22,27,42^. Cells carrying 2-stage and even 4-stage parasites, of which the latter have comparable sizes to the host nucleus and fill the entire width of the microchannel (**Figures 1A** and **1B**), still showed rapid migratory velocities of around 10 micrometers per minute, comparable to the velocity of non-infected bystander DCs (**Figure 1C** and **Supplemental Movie S1**). When the parasites further replicated to 8-, 16-, and 32-stages, the velocity of the host cell decreased but still reached substantial speeds of 3-8 micrometers per minute (**Figure 1C** and **Supplemental Movie S1**). Yet, notably, at these stages, the parasitic cargo is larger than the host nucleus, the bulkiest organelle in most cell types (**Figure 1B**). Thus, even cells that carried extremely large 16- and 32-stage parasites, which reside within a parasitophorous vacuole (PV) that is (i) up to 5 times longer than the host nucleus, (ii) entirely filled the width of the microchannels, and (iii) occupied large parts of the host cytoplasm (**Figures 1A, 1B** and **Supplemental Movie S1**), still moved in the range of micrometers per minute and thus a magnitude faster than motile mesenchymal cells.

To generalize these findings, we investigated the transport of large parasitic cargoes through constrained microenvironments by macrophages, an immune cell type that, in contrast to DCs, uses a slower adhesive mesenchymal migration strategy^10^. Notably, macrophages are preferentially infected by *Toxoplasma gondii*^43^ and are modulated by the parasite into a low-adhesive and highly motile state^25,26^. When we loaded macrophages upon their timed infection with *Toxoplasma gondii* parasites into linear microchannels (**Figure S1A**), quantification of their velocity revealed an extremely rapid motility of around 16 to 18 micrometers per minute of cells infected with 1-, 2-, and 4-stage parasites (**Figures 1D** and **S1A,** and **Supplemental Movie S2**). Comparable to infected dendritic cells, macrophages loaded with large 8-, 16- and 32-stage parasites carried parasitic cargoes that were up to 3 times larger than the macrophage’s nucleus (**Figures S1A and S1B**), slowed down but migrated still with substantial speeds of 3 to 10 micrometers per minute (**Figure 1D** and **S1A,** and **Supplemental Movie S2**). Importantly, these velocities represent rapid speeds for macrophages since they typically only migrate with speeds of around 1 micrometer per minute in their uninfected state^44^. Overall, these findings demonstrate that the transport of extremely large parasitic cargoes by immune cells through constrained microenvironments is remarkably efficient.

To identify the limits of parasitic transport by motile immune cells, we next introduced a defined pore of 4 micrometers in width into the migratory track (**Figure S1C**). Although larger parasitic cargoes took longer to squeeze through the pore within dendritic cells, they still passed surprisingly quickly (**Figures 1F** and **S1C**). To provide a smaller pore that is even smaller than the diameter of an individual parasite within the parasitophorous vacuole, we challenged parasite-infected DCs with pores of only 2 micrometers in width (**Figure 1E**). To our surprise, we still observed the rapid translocation of the host cell and its parasitic cargo through these narrow microenvironmental barriers (**Figures 1E** and **1G**, and **Supplemental Movie S3**). Dendritic cells infected with 1-, 2-, and 4-stage parasites took the same time to pass through the 2 micrometers pore as uninfected cells, and even large parasitic 16-stage cargoes only slowed down two-fold (**Figure 1G**). Similarly, parasite-infected macrophages crossed these narrow microenvironmental barriers with only a 2- to 3-fold delay at 4 to 8 and 16 to 32 parasite stages, respectively, compared to 1- to 2-staged parasites (**Figure S1D** and **Supplemental Movie S4**). Thus, *Toxoplasma* parasites efficiently overcome microenvironmental constraints and cross local barriers within infected dendritic cells and macrophages, even when the parasitic cargo reaches sizes substantially larger than any host cell’s organelle.

### Intracellular Parasites Cause Migration Bottlenecks for Infected Immune Cell Hosts

One solution to the paradox of how large parasitic cargoes might be transported efficiently through space-constrained microenvironments is that the parasite may be mechanically soft, enabling large deformations. To test this hypothesis, we performed atomic force microscopy (AFM) on *Toxoplasma*-carrying DCs after fixation in removable microenvironments that confine cells between two surfaces^45^ (**Figure 2A**). This approach allowed us (i) to measure the physical properties of the parasites and the host cells from the outside when they are flatly spread, and (ii) to uncouple the measurement of stiffness from the motility of the host cell. Quantification of the Young’s modulus, a mechanical property that increases with increasing stiffness, showed a higher Young’s modulus of each individual parasite in comparison to the local intra-vacuolar matrix between the individual parasites within a parasitophorous vacuole (PV) (**Figures 2A** and **2B**). Surprisingly, the stiffness of the parasites was even slightly higher than that of the host nucleus (**Figure 2A** and **2B**), which typically represents the major deformation bottleneck for motile cells crossing constrained microenvironments^2,4,5^. When we analyzed the dissipated viscous energy measured by AFM as a mechanical parameter for viscosity^46,47^, we detected a slightly lower dissipated viscous energy of the parasites compared to the host nucleus (**Figure 2A** and **2C**), indicating a more pronounced elastic response of the parasite. These data suggest that *Toxoplasma*-infected immune cells harbour a novel, stiff yet elastic, intracellular obstacle in addition to the bulky host nucleus.

**Figure 2.**
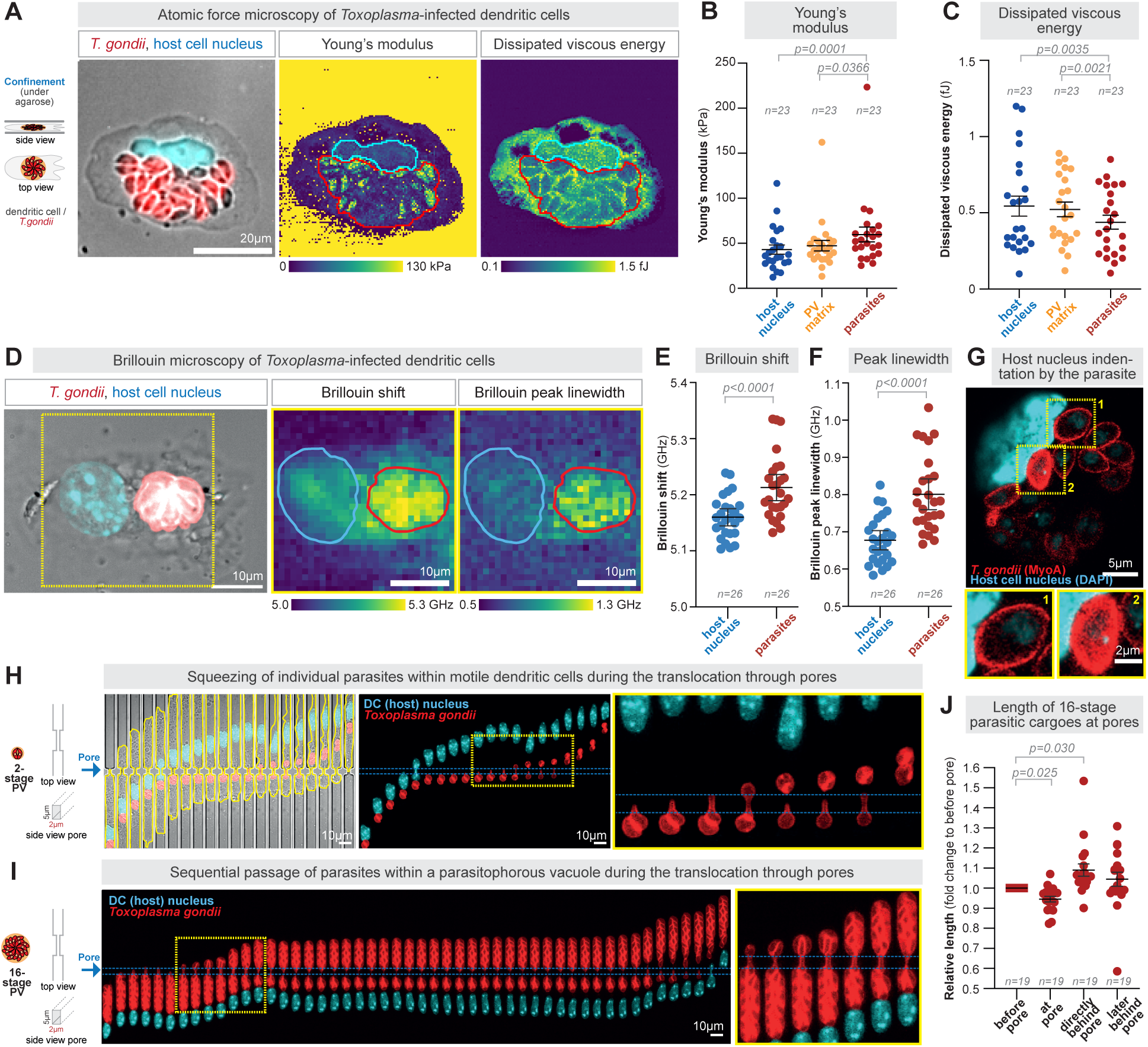
Intracellular parasites cause migration bottlenecks for infected immune cell hosts. **(A)** Representative fluorescent microscopy (left panel, T. gondii in red, host nucleus in blue) and atomic force microscopy (Young’s modulus, middle panel; dissipated viscous energy, right panel) of a migrating dendritic cell in confinement and upon PFA-based fixation. **(B)** Quantification of A), quantifying the Young’s modulus of the dendritic cell’s nucleus (host cell), the individual parasites, and the regions within the parasitophorous vacuole (PV) between the individual parasites (“PV matrix”). N = 23 cells from 4 independent biological replicates. Data are the mean±SEM. Statistics: Friedman ANOVA. **(C)** Quantification of A), quantifying the dissipated viscous energy of the dendritic cell’s nucleus (host cell), the individual parasites, and the PV matrix. N = 23 cells from 4 independent biological replicates. Data are the mean±SEM. Statistics: Friedman ANOVA. **(D)** Representative simultaneous fluorescent (left panel) and Brillouin microscopy (Brillouin shift, middle panel; Brillouin linewidth, right panel) images of a *Toxoplasma*-infected dendritic cell migrating in confinement and upon PFA-based fixation. **(E)** Quantification of D), quantifying the Brillouin shift of intracellular *Toxoplasma* parasites in comparison to the dendritic cell’s nucleus. N = 26 cells from 4 independent biological replicates. Data are mean±95% CI. Statistics: Wilcoxon matched-pairs signed rank test. **(F)** Quantification of D), quantifying the Brillouin peak of intracellular *Toxoplasma* parasites in comparison to the dendritic cell’s nucleus. N = 26 cells from 4 independent biological replicates. Data are the mean±95% CI. Statistics: Wilcoxon matched-pairs signed rank test. **(G)** Representative indentation of the host nucleus by *Toxoplasma* parasites. See Figure S2A for quantification. **(H)** Representative example of the squeezing of individual parasites in a 2-stage parasitophorous vacuole (PV) within a dendritic cell during passage through a 2 micrometer-sized pore. The yellow dashed square is shown in a zoom. **(I)** Representative 16-stage example showing unfolding of the parasitophorous vacuole (PV) during passage within a dendritic cell through a 2 micrometer-sized pore. The yellow dashed square is shown in a zoom. **(J)** Quantification of the change in length of 16-stage parasites during passage through 2 micrometer-sized pores. N= 19 cells from 3 independent biological replicates. Data are the mean±SEM. Statistics: Friedman ANOVA.

To corroborate this finding, we measured the internal mechanical properties of the parasites within infected immune cells using Brillouin microscopy^48^. Combining this approach with confocal imaging of the host cell nucleus and SAG1-Halo-labeled *Toxoplasma* parasites (**Figure 2D**) revealed a higher Brillouin shift (**Figure 2D** and **2E**) and a higher Brillouin peak linewidth (**Figure 2D** and **2F**) of the entire parasitic cargo in comparison to the host cell’s nucleus. The increased Brillouin peak linewidth, a proxy for the viscosity of the sample^48^, suggests that rather the low viscosity of the individual parasites measured with AFM is relevant during squeezing than the internal property of the entire cargo measured by Brillouin microscopy. The increased Brillouin shift, a proxy for the longitudinal modules and thus stiffness, further validates that the entire parasitic cargo is stiffer than the host nucleus. Together, these data support the finding that *Toxoplasma* parasites within motile immune cells have physical properties that make their deformation challenging.

Indeed, when we measured the shape of the parasite and the host nucleus when they were in direct neighborhood, we observed strong deformations of the host nucleus by individual parasites (**Figures 2G** and **S2A**), confirming a lower deformability of the parasite in comparison to the stiffest host organelle. We further validated this finding with live-cell imaging, which revealed dynamic and frequent nuclear deformation by the parasitic cargo when they were in close proximity (**Figure S2B**). Thus, intracellular *Toxoplasma* parasites are not only a large but also a physically challenging cargo.

To directly test whether this physically challenging cargo represents a new primary bottleneck for infected cells to move through constrained microenvironments, we next performed fast time-lapse and high-resolution imaging of labeled parasites translocating within either dendritic cells or macrophages through 2 micrometer pores (**Figures 2H** and **2I**, and **Supplemental Movie S3**). This revealed that large parasitic cargoes are compressed once they encounter the pore (**Figure 2J** and **Supplemental Movie S3**), followed by the sequential passage of individual parasites through the pore (**Figures 2H, 2I, S2D,** and **S2E,** and **Supplemental Movies S3 and S4**). During this sequential passage of individual parasites, their ellipsoid shape was substantially deformed (**Figures 2H, 2I, S2D,** and **S2E**). Behind the pore, the parasites rapidly returned to their original shape and reorganized within their parasitophorous vacuole (**Figures 2H-J**). Notably, not only were large parasitic cargoes compressed in front of the pore (**Figure 2J**) but also the individual parasites frequently resided for some time at the entrance of the pore (**Figure S2C-E**), which was followed by a quick translocation through the pore once the individual parasite was deformed (**Figures 2H** and **2J**), both underscoring the physical challenge of the parasitic cargo for passage through constrained microenvironments. Indeed, when we measured the time for the parasitic cargo to pass through the pore, compared with the passage times of the host nucleus and the entire host cell, we observed that the major translocation time is determined by large parasitic cargoes (**Figures S2F-I**), identifying that parasite-infected immune cells carry a novel intracellular bottleneck that is even more challenging than the bulky host nucleus. In line with these findings, we often observed an increased distance between the host nucleus and the parasitic cargo after the host nucleus passed through a constriction, while the parasitic cargo remained at the pore entrance (**Figure 2H and S2D**). Together, these results demonstrate that intracellular *Toxoplasma* parasites within motile immune cells are large, bulky, and rigid cargoes that locally unfold and re-fold to allow single parasites to translocate sequentially. Overall, these findings identify the parasitic cargo as the primary bottleneck during the migration of infected immune cells through constrained microenvironments.

### Intracellular Positioning Adapts to the Size of the Parasitic Cargo

To identify the mechanism by which infected immune cells migrate in constrained environments while carrying the physical burden of their parasitic cargo, we first quantified the intracellular position of the parasite relative to the host nucleus, as the position of organelles serves as an indicator of underlying cell polarity and thus cytoskeletal forces^49,50^. To our surprise, we observed that parasitic cargoes are non-randomly positioned within the cytoplasm of motile DCs, depending on the parasitic cargo stage (**Figures 3A** and **3B**, and **Supplemental Movie S1**). While small 1-stage parasitic cargoes are mostly positioned in the back of the host nucleus, growing parasitic cargoes are positioned with increasing frequency towards the cellular leading edge in front of the host nucleus, until very large 32-stage PV cargoes are predominantly positioned towards the host cell front (**Figure 3B**). To confirm this re-localization of the parasitic cargo with increasing size, we analyzed parasite-infected DCs in microchannels of larger width (**Figure S3A** and **S3B**) as well as during their navigation through maze-like microenvironments (**Figure S3C**). In all microenvironments, large parasitic cargoes were particularly frequently positioned in front of the host nucleus (**Figures 3B, S3A-C**). This even more forward positioning of large parasitic cargoes compared to the host nucleus is particularly unexpected, since DCs belong to the class of migrating cells that employ an amoeboid migration mode that is characterized by low-adhesiveness to the substrate and forward positioning of the nucleus towards the cellular protrusion^49,51^. Overall, these data reveal that the *Toxoplasma* parasitic cargo is positioned in a manner that adapts to its size.

**Figure 3.**
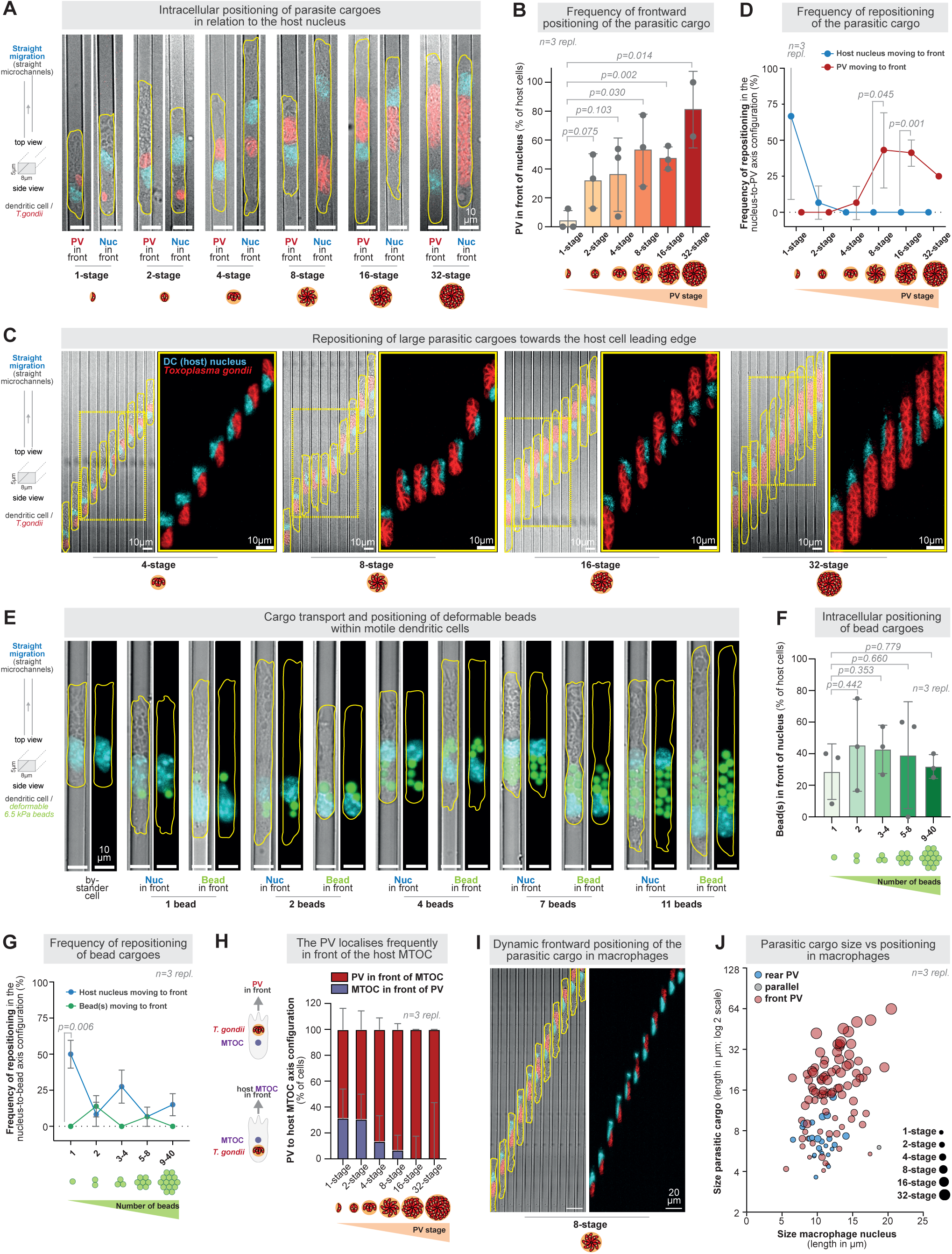
Intracellular positioning adapts to the size of the parasitic cargo. **(A)** Representative examples of the intracellular positioning of parasites forward or rearward relative to the host cell nucleus during dendritic cell migration in a straight 8 micrometer-wide microchannel. *Toxoplasma gondii* in red, host cell nucleus in cyan, host cell shape in yellow. **(B)** Quantification of the positioning of the parasite cargo in relation to the dendritic cell’s host nucleus, as shown in B). Note the increasing frequency of parasitic frontward positioning with its size. Data are the mean±SD. Unpaired t-tests. N=3 individual biological replicates (23 cells 1-stage PV, 29 cells 2-stage PV, 52 cells 4-stage PV, 99 cells 8-stage PV, 76 cells 16-stage PV, and 12 cells 32-stage PV). **(C)** Representative examples of dynamic repositioning of PV cargoes in front of the host cell nucleus towards the host cell front. **(D)** Quantification of dynamic repositioning events in the parasite to host nucleus axis, as shown in C). Note the increasing frequency of parasitic forward positioning with its size. Data are the mean±SD. Unpaired t-tests. N=3 individual biological replicates (23 cells 1-stage PV, 29 cells 2-stage PV, 52 cells 4-stage PV, 99 cells 8-stage PV, 76 cells 16-stage PV, and 12 cells 32-stage PV). **(E)** Representative dendritic cells with deformable (6.5 kPa) beads as cargoes, either positioned forward or rearward of the dendritic cell’s nucleus. Note how multiple beads cluster, comparable to the parasitic assemblies. Deformable beads in green, dendritic cell nucleus in cyan, dendritic cell shape in yellow. **(F)** Quantification of the intracellular positioning of deformable beads relative to the host cell nucleus, as shown in E). Data are the mean±SD. Unpaired t-tests. N=3 individual biological replicates (44 cells w/o beads, 25 cells 1 bead, 19 cells 2 beads, 31 cells 3-4 beads, 21 cells 5-8 beads, 22 cells 9-40 beads). **(G)** Quantification of dynamic repositioning events in the deformable bead to the dendritic cell’s nucleus axis. Data are the mean±SEM. Unpaired t-tests. N=3 individual biological replicates (44 cells w/o beads, 25 cells 1 bead, 19 cells 2 beads, 31 cells 3-4 beads, 21 cells 5-8 beads, 22 cells 9-40 beads). **(H)** Quantification of the positioning of the parasite in relation to the host cell’s microtubule-organizing center (MTOC) marked by EB3-mCherry, as shown in Figure S4E. Data are the mean±95% CI. N=3 individual biological replicates (19 cells 1-stage PV, 26 cells 2-stage PV, 22 cells 4-stage PV, 44 cells 8-stage PV, 18 cells 16-stage PV, and 5 cells 32-stage PV). See also the quantification of the parasite positioning in relation to the nucleus and the MTOC, as shown in Figure S4F. **(I)** Representative macrophage carrying an 8-stage parasite, in which the parasitic cargo dynamically repositions forward of the macrophage’s nucleus. **(J)** Quantification of the positioning of the parasite cargo in relation to the macrophage’s host nucleus. See also representative examples in Figure S1A. Data are the mean±SD. N=3 individual biological replicates (109 cells).

In agreement with size-dependent cargo positioning, we frequently observe the repositioning of large parasitic cargoes from the rear to the front during our imaging intervals but rarely vice versa (**Figures 3C, 3D**, **S3D** and **S3E,** and **Supplemental Movie S5**). Since these repositioning events of the large parasitic cargo require overtaking of the nucleus and thus substantial deformation of both cargoes (**Figure 3C, S3D** and **Supplemental Movie S5**), these results demonstrate that parasitic cargoes actively adapt their intracellular localization.

To confirm that repositioning is an active process that is specific to the parasitic cargo, we loaded parasite-infected DCs with rigid polystyrene beads. With a diameter of 6 micrometers, these beads mimic the size of a 4-stage parasitic cargo. However, we observed the beads mostly positioned behind the parasitic cargo (**Figures S4A** and **S4B**), suggesting that a passive bead cargo differs from a parasitic cargo. To further corroborate this finding, we loaded deformable beads (acrylamide co-acrylic acid microparticles^52,53^) with a size of 6 micrometers and a stiffness of 6.5 kPa into motile DCs. This approach has the advantages of mimicking (i) the size and (ii) the stiffness of individual parasites, also when compared to 1-5 kPa stiffness range of immune cell nuclei^54^. Indeed, as for parasitic cargoes, these bead cargoes dynamically deform the host cell nucleus (**Figure S4C** and **Supplemental Movie S6**). Moreover, DCs frequently took up multiple beads that clustered intracellularly, allowing quantification of cells carrying a range of bead assemblies and thus cargo sizes (**Figure 3E**), thereby closely mimicking the intracellular sizes of parasitic cargoes. However, in contrast to parasitic cargoes, these bead cargoes often localized to the rear half of the host cell body and randomly in relation to the host nucleus (**Figure 3F** and **3G**). Moreover, bead cargoes did not show a correlation between their positioning and cargo size (**Figure 3F**). Additionally, in contrast to parasitic cargoes, the intracellular position of the bead cargo remained stable along long migration tracks due to rare reposition events of the bead cargoes to the front of the host nucleus (**Figures 3G** and **S4D,** and **Supplemental Movie S6**). Together, these findings reveal that the cellular organization and behavior differ depending on whether the cells carry large bead-or parasitic-cargoes.

To characterize the adaptation of cellular organization to the size of the parasitic cargo, we next mapped the position of the parasite in relation to the microtubule-organizing center (MTOC), a key marker of cellular organization^55^. To do so, we infected EB3-mcherry expressing DCs, which serves as a microtubule-plus end and microtubule-organizing center (MTOC) marker within the host cell^12,56,57^. This revealed that large parasitic cargoes are exclusively positioned forward of the MTOC (**Figures 3H** and **S3E,** and **Supplemental Movie S7**), which typically localizes approximately to the cell center^49,55^. Moreover, in most cases, the large parasitic cargo is even positioned together with the nucleus in front of the MTOC (**Figure S3F,** and **Supplemental Movie S7**). These data show that both the parasitic cargo and the nucleus can simultaneously acquire an amoeboid-like positioning in front of the MTOC, and that the parasitic cargo preferentially locates forward of these key host organelles, particularly when it is large.

To test whether large parasitic *Toxoplasma* cargoes generally localize towards the cell front in infected immune cells, we infected macrophages and again observed parasitic cargoes actively positioned in front of the host nucleus (**Figures 3I**) in a size-dependent manner (**Figures 3J**), comparable to parasitic cargoes in motile DCs. Together, these results reveal that *Toxoplasma* parasitic cargoes actively position towards the front of their immune host cells despite their large, bulky nature. Overall, these findings reveal a mechanical adaptation of forces and intracellular positioning in response to the size of the parasitic cargo.

### Myosin-driven Host Cell Contractility Adapts to the Parasite Size

To identify the underlying mechanism of size-adaptive parasite positioning within infected immune cells, we developed a particle-based model to simulate the arrangement of the parasitic cargo relative to the host nucleus during immune cell migration. By combining dissipative particle dynamics^58^, with a representation of the host nucleus, the host cortex, and the parasitic cargo as elastic shells, this simulation captures the effect of (i) mechanical coupling between the host nucleus, the host cortex and the extracellular microenvironment, (ii) cytoplasmic flows, (iii) cell contractility, and (iv) parasite cargo size onto the localization of intracellular cargoes in a spatiotemporal manner (see Supplementary Information for details on the simulation). The position of the parasitic cargo is determined by a balance of pressure at the rear, representing actomyosin activity, and visco-elastic forces that result from a coupling between the cortex and the cargo. The simulation predicts that the pushing and pulling forces on the cargo depend on its size in constricted environments. Moreover, they revealed that increasing the host contractility, simulated by the host cortical tension, results in a higher likelihood of forward localization of the parasitic cargo, in particular when the parasitic cargo reaches sizes larger than the host nucleus (**Figures 4A** and **4B**, and **Supplemental Movie S8**).

**Figure 4.**
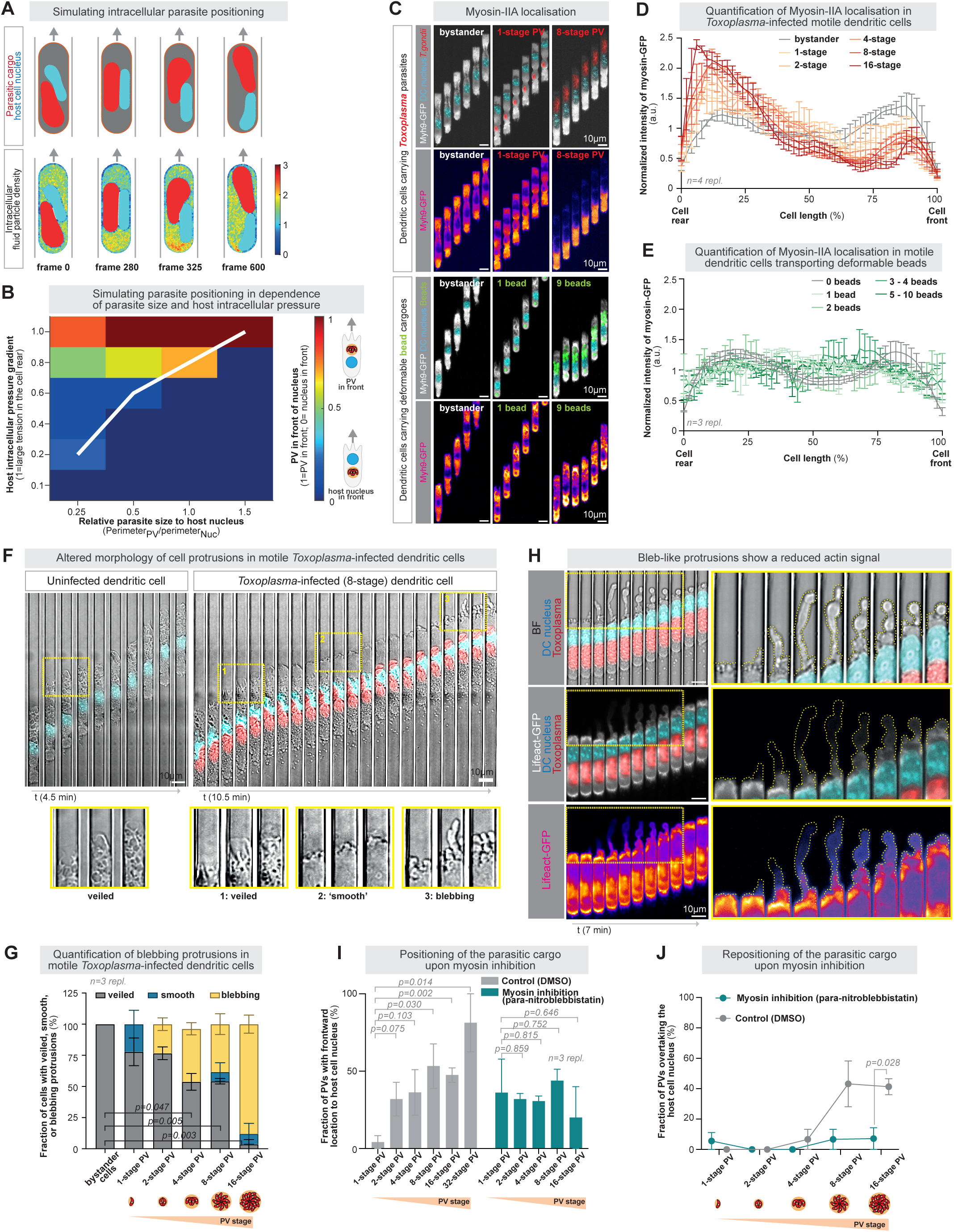
Myosin-driven host cell contractility adapts to parasite sizes. **(A)** Representative snapshots of simulating the intracellular positioning of parasites and the host nucleus in moving cells. *Toxoplasma gondii* in red, host cell nucleus in cyan, the host cytoplasm in gray (upper panel), the cell cortex in orange (upper panel), and the intracellular fluid particle density of the host in fire-color (lower panel). Note the repositioning of the parasite frontward of the host nucleus. **(B)** Simulation of parasite positioning in dependence of parasite size relative to the host nucleus, and the tension in the host cell rear that leads to an intracellular pressure gradient. **(C)** Representative examples of *Toxoplasma*-infected (upper panel) or bead-loaded (lower panel) Myh9-GFP (Myosin-II)-expressing dendritic cells migrating along linear paths. *Toxoplasma gondii* in red, host cell nucleus in cyan, Myh9-GFP in white or as fire-color. **(D)** Quantification of the Myh9-GFP signal over the cell length in *Toxoplasma*-infected dendritic cells. N=4 individual biological replicates (50 bystander cells, 11 cells 1-stage PV, 15 cells 2-stage PV, 38 cells 4-stage PV, 57 cells 8-stage PV, 30 cells 16-stage PV, and 4 cells 32-stage PV). **(E)** Quantification of the Myh9-GFP signal over the cell length in dendritic cells loaded with deformable beads. N=3 individual biological replicates (83 cells w/o beads, 73 cells 1 bead, 40 cells 2 beads, 37 cells 3-4 beads, 26 cells 5-10 beads). **(F)** Representative examples of protrusion shapes in uninfected or *Toxoplasma*-infected dendritic cells. Note the frequent appearance of smoother or even blebbing protrusions in infected dendritic cells, while uninfected cells typically show classical veil-like protrusions. *Toxoplasma gondii* in red and host cell nucleus in cyan. **(G)** Quantification of the frequency of veiled, smooth, or blebbing protrusions in uninfected or *Toxoplasma*-infected dendritic cells, as shown in F). Data are the mean±SEM. One-way ANOVA with Geisser-Greenhouse correction on the fraction of cells with veils. N=3 individual biological replicates (28 bystander cells, 12 cells 1-stage PV, 14 cells 2-stage PV, 14 cells 4-stage PV, 20 cells 8-stage PV, and 23 cells 16-stage PV). **(H)** Representative example of Lifeact-GFP-expressing, *Toxoplasma*-infected dendritic cell. Zoom-ins in the yellow boxes highlight a reduced actin signal within bleb-like protrusions, which is a typical feature of blebs. See also Figure S5A for comparison of uninfected bystander cells. **(I, J)** Positioning (I) and repositioning (J) of the parasitic cargo in the presence of the myosin II inhibitor para-nitro-blebbistatin (pNB) or DMSO controls. Note that the control cells are the same as in Fig. 3b. N=3 individual biological replicates (I: 22 cells 1-stage PV DMSO, 16 cells 1-stage PV pNB, 29 cells 2-stage PV DMSO, 34 cells 2-stage PV pNB, 50 cells 4-stage PV DMSO, 55 cells 4-stage PV pNB, 93 cells 8-stage PV DMSO, 62 cells 8-stage PV pNB, 67 cells 16-stage PV DMSO, 20 cells 16-stage PV pNB; J: 17 cells 1-stage PV DMSO, 8 cells 1-stage PV pNB, 18 cells 2-stage PV DMSO, 22 cells 2-stage PV pNB, 34 cells 4-stage PV DMSO, 38 cells 4-stage PV pNB, 69 cells 8-stage PV DMSO, 35 cells 8-stage PV pNB, 16 cells 16-stage PV DMSO, 16 cells 16-stage PV pNB). Unpaired t-tests. Data are the mean±SEM.

To directly test the prediction of the model experimentally, we imaged the localization of host myosin-IIA, since myosin-IIA is a key driver of cellular contractility in immune cells^36^, and since the myosin-II upstream regulators Rho/ROCK are critical for the tissue dissemination of the parasite by monocytes and macrophages^30^. Live cell imaging of non-muscle myosin heavy chain IIA (myosin-IIA, encoded by the MYH9 gene) using DCs encoding MYH9-GFP and infected with *Toxoplasma* parasites revealed a strong enrichment of myosin-IIA in the rear of the host cell behind the parasitic cargo (**Figure 4C** and **Supplemental Movie S9**). While uninfected immature DCs also often exhibit a rearward myosin accumulation^59^, the intracellular distribution pattern was more evenly distributed in uninfected bystander DCs when compared to parasite-infected DCs (**Figure 4C** and **D**). Notably, the more the parasite size increased, the more prominently myosin-IIA was enriched to the rear (**Figure 4D**). This finding demonstrates that large parasitic cargoes cause a shift in the distribution of host myosin-IIA from the front to the rear, which is indicative of rapid immune cell motility^59^. In contrast, DCs loaded with bead cargoes of different sizes did not show a correlation between cargo size and myosin-IIA enrichment at the host cell rear (**Figures 4C** and **4E**). Thus, the distribution of host myosin-IIA adapts to the size of the parasitic cargo.

To test whether redistribution of host myosin-IIA alters cellular contractility, we first examined the morphology of DCs infected with parasites. While motile DCs are morphologically characterized by veil-like protrusions and many macropinosome-like vesicles in the front part of the cell body^60,61^, *Toxoplasma gondii*-infected DCs frequently showed roundish, smooth, and vesicle-free cell fronts that can even have bleb-like morphologies (**Figure 4F** and **Supplemental Movie S10**), both indicators of a high degree of intracellular contractility. Notably, when we quantified the frequency of this smooth cell front phenotype in relation to the size of the parasitic cargo, we observed that large parasitic cargoes almost always caused the appearance of a non-veiled, smooth, or blebbing cell front (**Figure 4G**). A key defining feature of a bleb during bleb-based migration strategies is a high contractility, leading to the expansion of the bleb faster than the polymerization of F-actin^62,63^. By infecting DCs that express Lifeact-GFP as a marker for F-actin, we indeed observed rapidly expanding blebs that were devoid of F-actin (**Figure 4H** and **S5A,** and **Supplemental Movie S10**). Thus, large parasitic cargoes induce high contractility within their motile host immune cells. To further corroborate this finding, we depolymerized the host microtubule cytoskeleton with low doses of nocodazole, well-known to increase cellular contractility by the release of microtubule-bound myosin activators like RhoGEFH1/Lfc1^57,64^, and observed that the blebbing phenotype already appeared even with smaller parasitic cargoes (**Figure S5B** and **S5C**). Together, these data identify an enhanced myosin-based contractility in cells carrying large parasitic cargoes.

To directly test whether this enhanced myosin-based contractility causes the forward positioning of large parasitic cargoes, as predicted by our simulations, we used the myosin inhibitor para-nitroblebbistatin and measured the localization of the parasitic cargo during DC migration. While control cells positioned their large parasite cargoes forward, myosin-inhibited cells displayed a localization of the parasitic cargo independent of its size (**Figure 4I**). This finding was substantiated when we quantified the frequency of repositioning of the parasitic cargo and the host nucleus along long migration tracks, showing that myosin inhibition strongly impaired the ability of parasitic cargoes to move towards the host cell front (**Figure 4J**). Thus, myosin-based contractility within the host cell drives the intracellular positioning of the parasite as an adaptation to its size. Overall, these findings reveal a shift in the force balance of motile immune cells hijacked by *Toxoplasma* parasite, depending on the pathogen’s size.

### Adaptive Myosin-driven Contractility Facilitates Parasitic Transport Across Local Barriers

These findings suggest a model in which myosin-based contractility at the host cell’s rear generates a contractile force that propels the parasitic cargo through space-limiting environments in a size-dependent manner. To experimentally test this model, we infected DCs and measured the force they generate in their local microenvironment using quantitative traction force microscopy (TFM). Since motile immune cells, and particularly *Toxoplasma-*infected immune cells, use an amoeboid migration strategy that is non-adhesive and therefore prevents their analysis on flat two-dimensional substrates^10,21^, we measured the traction forces generated by cells confined between two substrates, one of which had embedded fluorescent beads (**Figure 5A** and **Supplemental Movie S11**). Mapping and quantification of the displacement of the beads during the motility of infected DCs (**Figure S5D**) showed higher bead displacements (**Figure S5E**) and, therefore, higher traction magnitudes (**Figure 5B**) of cells carrying large parasitic cargoes. These results show that immune cells can adjust their force-regime to the size of the parasitic cargo.

**Figure 5.**
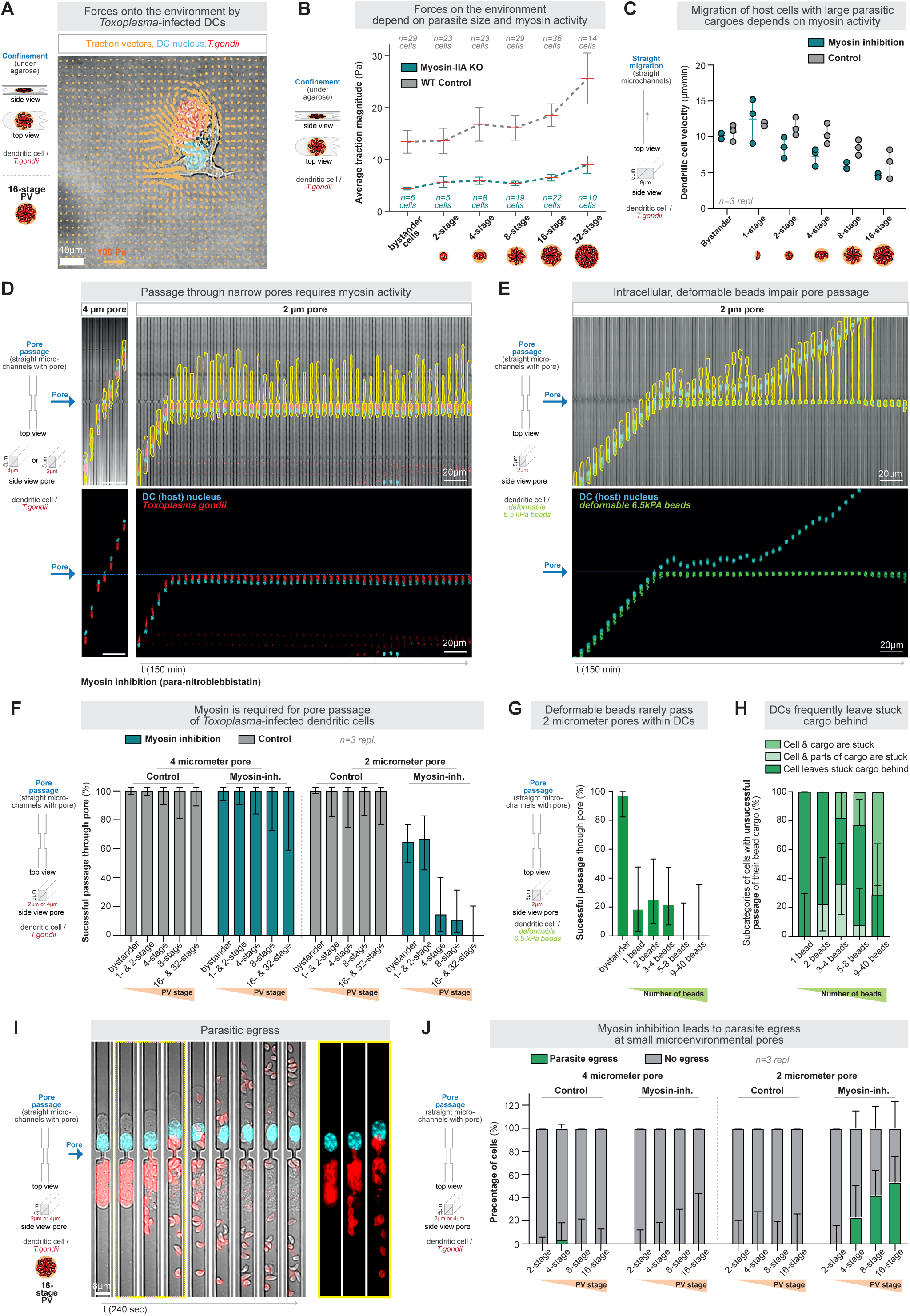
Adaptive myosin-driven contractility facilitates transport across local barriers. **(A)** Representative traction force measurement of a dendritic cell infected with *Toxoplasma gondii* (16-stage PV) migrating in confinement. Traction vectors are shown in orange, the DC nucleus in cyan, and the parasite in red. See Supplemental Figure 5C for representative examples of bead displacements that are the basis for the traction vector calculation, and Supplemental Figure 5D for the quantification of bead displacements. **(B)** Quantification of A), showing the average traction magnitude (Pa) of WT control cells and myosin-IIA knockout cells infected with *Toxoplasma gondii*. Note the increase in traction magnitudes in WT cells with increasing parasitic sizes, and the reduced traction magnitudes in myosin-IIA knockout cells. N = 29 cells (1-stage PV, Control), 23 cells (2-stage PV, Control), 23 cells (4-stage PV, Control), 29 cells (8-stage PV, Control), 36 cells (16-stage PV, Control), and 14 cells (32-stage PV, Control), and 6 cells (1-stage PV, Myosin-IIA KO), 5 cells (2-stage PV, Myosin-IIA KO), 8 cells (4-stage PV, Myosin-IIA KO), 19 cells (8-stage PV, Myosin-IIA KO), 22 cells (16-stage PV, Myosin-IIA KO), and 10 cells (32-stage PV, Myosin-IIA KO) from 5 independent biological replicates. Data are the mean±SEM. **(C)** Dendritic cell migration velocity through 8 micrometer-wide microchannels upon *Toxoplasma gondii* infection, in the presence of the myosin II inhibitor para-nitroblebbistatin (pNB) or DMSO controls. N=3 individual biological replicates (86 cells (bystander, Control), 23 cells (1-stage PV, Control), 29 cells (2-stage PV, Control), 52 cells (4-stage PV, Control), 99 cells (8-stage PV, Control), and 76 cells (16-stage PV, Control), and 103 cells (bystander, pNB), 16 cells (1-stage PV, pNB), 34 cells (2-stage PV, pNB), 55 cells (4-stage PV, pNB), 62 cells (8-stage PV, pNB), and 20 cells (16-stage PV, pNB)). Data are the mean±SD. **(D)** Representative kymographs of *Toxoplasma gondii*-infected motile dendritic cells encountering 4 (left) and 2 (right) micrometer-wide microchannel pores in the presence of the myosin II inhibitor para-nitro-blebbistatin. *Toxoplasma gondii* is shown in red, the host cell shape in yellow, and the DC nucleus in cyan (Hoechst). **(E)** Representative kymographs of bead(deformable)-loaded motile dendritic cells encountering 2 micrometer-wide microchannel pores. Deformable beads in green, dendritic cell nucleus in cyan, dendritic cell shape in yellow. **(F)** Quantification of successful passage of dendritic cells through 4 and 2 micrometer-wide microchannel pores upon *Toxoplasma gondii* infection, in the presence of the myosin II inhibitor para-nitroblebbistatin (pNB) or DMSO controls. N=3 individual biological replicates (84 cells (bystander, 2 µm, Control), 15 cells (1-2-stage PV, 2 µm, Control), 10 cells (4-stage PV, 2 µm, Control), 16 cells (8-stage PV, 2 µm, Control), and 11 cells (16-32-stage PV, 2 µm, Control), and 48 cells (bystander, 2 µm, pNB), 21 cells (1-2-stage PV, 2 µm, pNB), 14 cells (4-stage PV, 2 µm, pNB), 19 cells (8-stage PV, 2 µm, pNB), and 15 cells (16-32-stage PV, 2 µm, pNB), and 74 cells (bystander, 4 µm, Control), 63 cells (1-2-stage PV, 4 µm, Control), 28 cells (4-stage PV, 4 µm, Control), 14 cells (8-stage PV, 4 µm, Control), and 26 cells (16-32-stage PV, 4 µm, Control), and 37 cells (bystander, 4 µm, pNB), 28 cells (1-2-stage PV, 4 µm, pNB), 17 cells (4-stage PV, 4 µm, pNB), 9 cells (8-stage PV, 4 µm, pNB), and 5 cells (16-32-stage PV, 4 µm, pNB)). Data are the fraction of the total±95% CI. **(G)** Quantification of successful passage of bead(deformable)-loaded dendritic cells through 2 micrometers-wide microchannel pores. N=3 individual biological replicates (28 cells w/o beads, 11 cells 1 bead, 12 cells 2 beads, 14 cells 3-4 beads, 13 cells 5-10 beads). Data are the fraction of the total±95% CI. **(H)** Quantification of the frequency of subcategories, if deformable-bead loaded dendritic cells do not successfully pass the 2 micron-sized pore. N=3 individual biological replicates (9 cells 1 bead, 9 cells 2 beads, 11 cells 3-4 beads, 13 cells 5-8 beads, 7 cells 9-40 beads). Data are the fraction of the total±95% CI. **(I)** Representative parasitic egress from a *Toxoplasma*-infected dendritic cell through a 2 micron-pore. **(J)** Quantification of parasitic egress upon migration or attempt migration through 4 and 2 micrometer-wide microchannel pores upon *Toxoplasma gondii* infection, in the presence of the myosin II inhibitor para-nitroblebbistatin (pNB) or DMSO controls. N=3 individual biological replicates (15 cells (1-2-stage PV, 2 µm, Control), 10 cells (4-stage PV, 2 µm, Control), 16 cells (8-stage PV, 2 µm, Control), and 11 cells (16-32-stage PV, 2 µm, Control), and 20 cells (1-2-stage PV, 2 µm, pNB), 13 cells (4-stage PV, 2 µm, pNB), 19 cells (8-stage PV, 2 µm, pNB), and 15 cells (16-32-stage PV, 2 µm, pNB), and 63 cells (1-2-stage PV, 4 µm, Control), 27 cells (4-stage PV, 4 µm, Control), 14 cells (8-stage PV, 4 µm, Control), and 26 cells (16-32-stage PV, 4 µm, Control), and 28 cells (1-2-stage PV, 4 µm, pNB), 17 cells (4-stage PV, 4 µm, pNB), 9 cells (8-stage PV, 4 µm, pNB), and 5 cells (16-32-stage PV, 4 µm, pNB)). Data are the fraction of the total±95% CI.

Next, to test whether this adaptive force generation is myosin-dependent, we infected DCs with a genetic knockout of myosin-IIA. This revealed that the overall average traction magnitude was lower in *Toxoplasma*-infected myosin-IIA DCs than in *Toxoplasma*-infected wild-type DCs, and increased less with the size of the parasitic cargo (**Figure 5B** and **S5E**). Together, these data demonstrate that immune cells adapt their force-generation to the size of the parasitic cargo in a myosin-IIA-dependent manner.

To test whether these myosin-IIA-dependent forces are required for parasitic cargo transport, we next quantified the velocity of parasite-loaded motile DCs upon myosin inhibition in linear 8-micrometer-wide microchannels. This demonstrated that myosin inhibition by para-nitroblebbistatin impaired the migration of DCs carrying larger parasitic cargoes (**Figure 5C**). In contrast, uninfected bystander cells and parasite-infected cells with small 1-stage PVs reached the same velocities upon myosin inhibition as control cells (**Figure 5C**). These data show that the migration of hijacked immune cells through confined microenvironments is facilitated by adaptive myosin contractility to the size of the parasite cargo.

To investigate the consequences for migration across local barriers, we next investigated how frequently parasite-infected DCs passed through 2 micrometers pores following myosin inhibition. While almost all control cells efficiently translocated through these small pores, myosin inhibition strongly impaired the translocation of larger PVs (**Figures 5D** and **5E**, and **Supplemental Movie S12**). While small 1-and 2-stage parasites successfully migrated through the pores in approximately 60% of cases, large 4-, 8-, and 16-stage parasites almost always got stuck at the 2 micrometer-sized pores (**Figures 5D** and **5E**). This behavior was reminiscent of cells loaded with assemblies of deformable beads, which rarely achieved the successful movement of the cargo across 2 micrometers pores (**Figures 5F-H**, and **Supplemental Movie S12**). Notably, when we imaged the behavior of parasite-loaded DCs that become stuck at the pore entrance, we observed parasite egress from the host (**Figure 5I** and **Supplemental Movie S12**). Quantification relative to the cargo size revealed that extracellular egress was rare in control cells but occurred frequently with large parasitic cargoes upon the impairment of myosin activity (**Figure 5J**). Together, these findings reveal that contractility adaptive to the size of the parasitic cargo is critical for the successful transport of *Toxoplasma* parasites within host cells through space-limited environments and across local barriers.

Overall, our results establish that the parasite *Toxoplasma gondii* alters the cellular force balance of its motile host cells by hijacking and adapting their myosin-based contractility. Thereby, we reveal a novel hijacking strategy in which the parasite modifies its host’s force regime and subcellular architecture to accommodate the size of its parasitic cargo. This mechano-adaptation to size facilitates the transport of large intracellular parasite assemblies across locally complex and confined microenvironments, and resolves the paradox of how parasite-infected immune cells can effectively migrate through such environments despite their bulky parasitic cargo.

## Discussion

Cell migration through confining tissue environments requires continuous coordination between cytoskeletal force generation, subcellular architecture, and the mechanical properties of intracellular organelles^1,2,9,36,49^. While the nucleus has emerged as a dominant mechanical bottleneck in confined migration^2,4,5^, cells like phagocytes can carry additional intracellular cargoes. However, whether such cargoes and their physical properties challenge the migratory capacity, and how cells respond to this challenge, has remained unknown. Here, we identify large intracellular parasites as previously unrecognised mechanical constraints for immune cell migration and uncover a mechanism by which host cells adaptively rewire cytoskeletal forces and intracellular organization to enable the transport of oversized, rigid cargoes through confining microenvironments.

Our findings reveal a novel principle of subcellular architecture dynamics and mechanoadaptation during cell migration: motile immune cells dynamically reorganize the spatial organization between force-generating cytoskeletal networks, the nuclear organelle bottleneck, and a mechanically demanding intracellular cargo. In uninfected cells, the nucleus typically represents the stiffest and largest intracellular object, having a defined intracellular position^50^ and acting as a dominant intracellular obstacle for migration through space-limiting environments^2,4,5,34,65,66^. In contrast, we show that intracellular *Toxoplasma gondii* parasites can exceed the nucleus in both stiffness and size, effectively overriding it as the primary mechanical obstacle. Remarkably, instead of preventing migration, infected cells reposition the parasitic cargo to the front of the host cell body as the size of the cargo burden increases, mediated by increasingly rearward-localized contractile forces. This adaptive positioning highlights the plasticity of subcellular architecture during migration. Notably, inert cargoes of comparable size and physical properties do not elicit this adaptive contractile response, in contrast to the previously discovered recruitment of the actin cytoskeleton to beads within confined migrating DCs^6^, suggesting that immune cells engage a cargo-selective or cargo-induced contractile program. More broadly, these findings establish that foreign cargoes influence immune cell trafficking, either by impeding passage through micron-scale pores, as observed for inert particles, or by inducing force and architecture rewiring, as observed for active parasitic cargoes.

From a mechanical perspective, our data demonstrate that immune cells adapt cytoskeletal force generation to the size of the parasitic cargo. We observe a progressive rearward localization of myosin-II and an increase in contractile, bleb-inducing forces as the parasite size increases, suggesting a scaling relationship between cargo dimensions and force output. The spatial distribution of myosin-II, the independence from microtubule-based pulling forces, the bleb-based migration mode, and the loss of efficient parasite translocation upon myosin inhibition together support a model in which rearward contractility predominantly generates pushing forces that drive the parasitic cargo through confinement. Notably, these mechanistic and mechanical findings are fully consistent with previous work showing that Rho/ROCK signaling, which is a key upstream regulator of myosin-II activity, is critical for *in vivo* tissue dissemination of *Toxoplasma* by monocytes and macrophages^30^.

In addition to force scaling, we uncover a mechanical shape-adaptation mechanism in which the parasitic cargo transiently deforms and unfolds into smaller substructures during the passage through narrow constrictions. This dynamic reduction in the effective cross-sectional area indicates that confined migration with oversized cargo is not determined solely by static size or stiffness, but also by the capacity for reversible cargo remodeling under force. This behavior mirrors principles observed at the cellular and nuclear levels, where cells exploit shape plasticity to traverse restrictive microenvironments^1^, and where neutrophil granulocytes use lobulated nuclear shapes to efficiently translocate through micron-scale constrictions^11,67^. Our findings extend these principles to intracellular cargo, highlighting that force-induced cargo remodeling represents an important layer of mechanical adaptation during migration.

Beyond cell biology and mechanics, these findings have important implications for parasite dissemination within host organisms. Intracellular pathogens are known to hijack host cytoskeletal forces to facilitate uptake, intracellular motility, stability, and evasion of degradation. For instance, bacterial pathogens such as *Shigella flexneri* exploit actin-driven macropinocytosis for entry^68^, *Chlamydia* relies on host actin scaffolds for vacuole stability^69^, and *Listeria* and *Rickettsia* hijack actin polymerisation for comet-like propulsion^70,71^. In contrast, how large eukaryotic intracellular parasites exploit host cytoskeletal forces has remained poorly understood. Our findings demonstrate that *Toxoplasma* parasites, unlike non-living bead cargoes of comparable sizes and stiffness, elicit a size-dependent reorganization of host contractility, indicating that the parasite is not a passive mechanical burden but an active modulator of host cell mechanics. This size-dependent force-rewiring of motile immune cells complements the known ability of *Toxoplasma* parasites to induce a rapid migratory, low-adhesive state in their immune cell hosts, together likely providing an evolutionary advantage by enabling a robust dissemination strategy across diverse tissues and physical environments.

In summary, our study uncovers a previously unrecognized intersection between subcellular architecture, adaptive force generation, and host-pathogen interactions. By revealing how migrating immune cells reorganize intracellular mechanics to accommodate oversized parasitic cargoes, we establish new principles of mechanoadaptation with implications for cell migration, organelle positioning, immune responses, and pathogen dissemination across tissues.

## Supporting information

Supplemental Information to the Computational Model

## Methods

### Cell culture

#### Dendritic cells, macrophages, and HoxB8-FL cells

Murine dendritic cells (DCs), macrophages, and immortalised hematopoietic precursor Hoxb8-FL cells^72^, were cultured at 37°C in a humidified incubator with 5% CO_2_. For cell culture, a basis medium composed of RPMI-1640 medium (Invitrogen, 21875034), 10% fetal bovine serum (Gibco, 10437-028), 1% penicillin-streptomycin (Sigma-Aldrich, P0781, 100 U/ml penicillin and 100 µg/ml streptomycin), and 50 µM 2-mercaptoethanol (Gibco, 31350-010), was supplemented with cell culture supernatant of growth-factor-expressing cells. DC medium was supplemented with 10% GM-CSF supernatant, macrophage medium was supplemented with 5% Flt3l supernatant and 20% M-CSF supernatant, and Hoxb8-FL medium was supplemented with 5% Flt3l supernatant and 1 µM estradiol (Sigma-Aldrich, E2758-1G). All centrifugations for washing or pelleting murine cells were performed at 300 g for 5 min. During experiments, all cells were maintained in imaging medium, phenol-free basis medium supplemented with 50 μM L-ascorbic acid (MilliporeSigma, W210901) to limit photobleaching and phototoxicity.

#### Bone marrow isolation from mice

All animal experiments were performed in accordance with the German Animal Welfare Act. The Core Facility Animal Models of the Biomedical Center (BMC) at LMU Munich housed the used mice. Bone marrow from male and female C57Bl6/J mice, sacrificed at 8 to 12 weeks of age, was used to generate bone marrow-derived dendritic cells (BMDCs). For this, mice were euthanised by cervical dislocation, femurs and tibiae were isolated, cut open and flushed with basis medium as previously described^73^.

#### Differentiation of dendritic cells and macrophages

DCs were differentiated from hematopoietic progenitor cells, either from murine bone marrow, obtained from male or female 8-to 12-week-old C57Bl6/J wild-type or MyoIIA-Flox*CD11c-Cre mice^12^, or from Lifeact-GFP, EB3-mCherry^57^, or Myh9-GFP Hoxb8-FL progenitor cell lines, as previously described^74^. In short, progenitor cells were seeded in DC medium. On day 3, 20% GM-CSF supernatant DC medium was added to the cells at a 1:1 ratio and on day 6, half the culture medium was replaced with 20% GM-CSF supernatant DC medium without pipetting the remaining cells. DCs were frozen on day 8, except for those with the MyoIIA-Flox*CD11c-Cre genetic modification, which were differentiated until day 10, with an extra medium change on day 8, and then frozen on day 10. DCs were frozen in fetal bovine serum (FBS, Gibco, 10437-028) supplemented with 10% DMSO, first for 3 days at-72 °C in a suitable freezing container, then stored in liquid nitrogen storage and used for experiments up to 1 year later.

Macrophages were differentiated from wild-type Hoxb8-FL progenitor cells, as described before^75^. Hoxb8-FL progenitor cells were cultured in macrophage medium, with medium changes on days 3 and 5, and macrophages were used for experiments on day 6 of the differentiation.

#### Culturing of Toxoplasma gondii parasites

*Toxoplasma gondii RH* Δ*ku80DiCre SAG1 gene* was tagged with Halo following standard protocols established in the lab ^31,41^. Parasites were maintained on HFFs (Human foreskin fibroblasts (HFFs) (RRID: CVCL_3285, ATCC) in DMEM (Dulbecco’s modified Eagle’s medium), 10% fetal bovine serum, 2 mM L[glutamine and 25 mg/ml gentamicin, and maintained at 37°C and 5% CO_2_. Cultured cells and parasites were screened against mycoplasma contamination with LookOut Mycoplasma detection kit (Sigma) and treated with Mycoplasma Removal Agent (Bio□Rad), if needed.

#### Infection of dendritic cells with Toxoplasma gondii parasites

HFFs 4 cm dish cell cultures were infected with 5×10^6^ parasites and left to replicate for 48 h until replicating vacuoles were ready to egress. Following the established protocols in the lab^76^, parasites were released by scratching and syringing, and then filtered to remove cell debris. Next, *Toxoplasma* cells were pelleted, resuspended in prewarmed PBS, washed again, and resuspended in prewarmed culture medium for the respective host cell type. For the infection with *Toxoplasma gondii,* differentiated DCs were thawed 2-4 h before infection in a water bath at 37 °C, washed with basis medium, resuspended in DC medium and incubated in a 6-well plate at 1×10^6^ DCs in 3 ml medium per well. For their infection, macrophages were grown close to confluency and used on day 6 of their differentiation after washing in PBS and adding 3 ml new macrophage medium per well of 6-well plates. DCs and macrophages were challenged with 0.5×10^6^ parasites per well for 20 h, constituting a ∼1:2 ratio of parasites to immune cells. Then, the cells were stained for 50 min with Janelia dye 646 HaloTag ligand (Promega, GA1120) at a 1:15000 dilution and two drops of NucBlue (Hoechst 33342, Invitrogen, R37605) were added per well without medium change. Infected macrophages lost adhesion and were harvested by flushing the culture dish surface with a 1 ml pipette. Stained DCs and MCs were pelleted and resuspended in imaging medium, optionally supplemented with inhibitors or DMSO for inhibitor experiments.

### Migration assays

#### PDMS microchannel assays

Microchannels were generated as previously described^39^. Polydimethylsiloxane (PDMS, Sylgard 184, Biesterfield) was mixed at a 1:10 elastomer-to-curing agent ratio, poured over a silicon wafer with negative microchannel structures imprinted by photolithography, degassed under vacuum, and cured at 80°C overnight. The PDMS was cut into blocks of approximately 7×12 mm. Two parallel holes, each 2×7 mm, were punched with a custom-made puncher at a 1 mm spacing in between. The devices and glass coverslips, cleaned with isopropanol and ethanol in an ultrasonic cleaning bath, were then cleaned in a plasma cleaner and bonded at 120°C. The PDMS devices were glued to a 6-well plate with 18 mm holes. After 2 min of treatment in the plasma cleaner, the microchannels were flooded with imaging medium and equilibrated in the cell culture incubator. Imaging medium supplemented with 0.625 µg/ml CCL19 was loaded into one hole, and 5×10^4^ *T. gondii*-infected and stained DCs were loaded into the second hole. Imaging was started after about 0.5-1 h, when the first cells entered the microchannels. Linear microchannels that were 8 μm wide and 5 μm high were used for migration analysis. For analysis of pore passage dynamics, the linear microchannels were constricted at a single point to 2 or 4 μm over a 5 μm length. For the analysis of rigid bead positioning within DCs, a pillar forest PDMS design was used, with a height of 5 μm, 7 μm-diameter pillars, and a 10 μm spacing between pillars.

#### Under-agarose migration assay

For cell migration under confinement in a 2D plane, DCs were allowed to migrate under 1% agarose towards the chemokine CCL19, as described before^45^. Briefly, agarose solution was prepared by mixing 1 part 2×HBSS, 2 parts phenol-free basis medium with 20% fetal bovine serum and 1 part high-molecular weight agarose (Biozym Gold Agarose, 850152) dissolved in pure water at 4% (w/v), adjusted to 55 °C, and the resulting 1% agarose mix was poured into 3.5 cm glass bottom dishes or per well of an 8-well slide, 3 ml or 300 μl, respectively. The agarose was solidified at room temperature for 1 h, then equilibrated in a cell culture incubator and in a petri dish containing a wet paper towel to provide additional humidity.

#### Inhibitor experiments

Inhibitors were added to the cells after the last washing step, directly after staining and before adding them to the experimental setup. Inhibitors were also added to the experimental environment and to the CCL19 medium. Para-Nitroblebbistatin was used at 25 µM, with the same 1:2000 DMSO dilution in the medium as a control. Nocodazole was used at 1 µM, with the same 1:3333 DMSO dilution as a control.

#### Deformable acrylamide co-acrylic acid beads

Deformable acrylamide co-acrylic acid beads with a diameter of 6.2 µm and a stiffness of 3.9 kPa were generated as previously described^52,53^. DAAM-particles (5% v/v) were covalently conjugated with 20 mg/mL BSA and 0.1 mM TMR-cadaverine (Invitrogen, A1318). First, they were washed twice (3000 g centrifugation for 1 min, and subsequent resuspension and mixing by vortexing) in activation buffer (100 mM MES pH 6, 200 mM NaCl). Particles were incubated in activation buffer with 0.1% (v/v) Tween 20, 40 mg/mL N-(3-Dimethylaminopropyl)-N’-ethylcarbodiimide hydrochloride, and 20 mg/mL N-hydroxysuccimide for 15 min while mixing. Particles were subsequently washed three times with PBS pH 6 with 0.2% (v/v) Tween 20. After the final wash, particles were resuspended in PBS pH 6 with 0.2% (v/v) Tween 20 at 10% (v/v) solids and mixed thoroughly. Then, 2x PBS pH 8.5 with 0.2% (v/v) Tween 20 was added and particles were directly incubated with BSA. After 1h, 0.1 mM TMR-cadaverine was added, and after another 0.5 h, blocking buffer consisting of 100 mM TRIS (pH 9), 100 mM NaCl and 100 mM ethanolamine was added. After 0.5 h, particles were washed three times in PBS pH 7.4 + 0.1% Tween 20. Beads were previously determined to have increased Young’s moduli of 6.5 kPa after BSA functionalization^52^. Beads were further opsonized with 0.03 mg/ml anti-BSA IgG antibody (MP Biomedicals, 08651111) in PBS with 0.1% Tween for 1 h under rotation, washed three times with PBS (pH 7.4) with 10% Tween with centrifugation steps at 3000 g for 1 min, and stored in PBS supplemented with 5 mM sodium azide. The beads were washed with PBS and resuspended in DC medium before adding them to DCs. Differentiated DCs were thawed and 3.3×10^5^ DCs were challenged with 1.5×10^6^ deformable beads for 20 h in 1 ml DC medium in wells of a 24-well plate. DCs were stained with one drop of NucBlue (Hoechst 33342, Invitrogen, R37605) per well for 20 min, and the medium was changed to imaging medium. Imaging during migration in linear microchannels or microchannels with 2 μm constrictions was performed at 1 frame per minute.

### Imaging

#### Live-cell imaging

Live-cell imaging was performed using a Leica DMi8 inverted wide-field epifluorescence microscope with a motorised stage, under cell culture conditions of 37°C and 5% CO_2_ in a humidified atmosphere (Pecon). Time-lapse recordings were performed using a 20x (Leica HC PL APO 20x/0.80, 11506529) or 40x (Leica HC PL APO 40x/0.95 CORR, 11506414) lens. The Leica sCMOS Camera DFC9000 GT was used as the camera, with LED5 fluorescence light source. *Toxoplasma* signal (Janelia dye 646) was imaged with a far-red filter (Excitation: 638/31, Emission: 695/58) and 640 nm excitation, EB3-mcherry signal with an orange filter (Excitation: 554/24, Emission: 594/32) and 550 nm excitation, and the nuclear signal (Hoechst 33342 or DAPI) with a blue filter (Excitation: 391/32, Emission: 435/30) and 395 nm excitation. Exposure to 395 nm light was kept to a minimum, just enough to clearly localize the nucleus position and size, to avoid phototoxicity affecting experimental results.

#### STED imaging

Fixation of cells was performed as described in Del Rosario et al.^76^. Briefly, cells were fixed in cytoskeleton buffer composed of two components: cytoskeleton buffer (CB1) (MES pH 6.1 10 mM, KCL 138 mM, MgCl 3 mM, EGTA 2 mM, 5% PFA) and cytoskeleton buffer (CB2) (MES pH6.1 10 mM, KCL 163.53 mM, MgCl 3.555 mM, EGTA 2.37 mM, sucrose 292 mM). These buffers were mixed in a 4:1 ratio, respectively, and subsequently used for fixation for 10 min. The samples were quenched (NH_4_CL 50 mM) for 10 min and washed three times in PBS. To visualise microtubules, rat anti-human alpha tubulin (BioRad) was used at 1:300 in 2% BSA 0.2% triton x100 in PBS for 1 h, washed three times in PBS, followed by an incubation with 1:1000 secondary antibody Alexa 488 (Life Technologies).

STED and confocal imaging were performed on an Abberior Expert Line STED microscope (Abberior Instruments GmbH, Göttingen, Germany) built around an Olympus IX83 inverted microscope body, equipped with an Olympus 100x / NA 1.4 Oil UPLSAPO objective and the easy3D STED module (spatial light modulator for 2D and 3D beam shaping). The system was controlled by Imspector software (Abberior Instruments). Samples were excited with 405 nm, and pulsed 485 nm, or 640 nm laser diodes at 1%, 2%, 5% of maximum power. Stimulated emission depletion was performed with a pulsed 595 nm STED laser at 5% of maximum power (repetition rate 40 MHz) using a 2D doughnut depletion pattern. The 405 nm channel was acquired only in confocal mode. Fluorescence was detected on single-photon-counting avalanche photodiodes (APDs) with the following spectral detection windows/ Filter Cubes: channel 1: 500–550 nm (485 nm excitation); channel 2: 650–720 nm (640 nm excitation). The pinhole was set to 1.0 Airy unit. Images were acquired with 40 nm pixel size, 4 µs pixel dwell time, and 10 line accumulations.

### Force and mechanical measurements

#### Traction force microscopy

Glass coverslips (Marienfeld, 60 × 24 mm, #1.5H) were cleaned in two stages. Initially, they were sonicated in a 5% v/v Hellmanex III solution for 10 minutes, followed by sonication in 100% ethanol. At each step, the coverslips were washed rigorously in double-distilled water for 5 minutes. Finally, the coverslips were dried using compressed nitrogen.

Silicone substrates were fabricated by combining a 1.2:1 weight ratio of two components, Cy-A and Cy-B (Dow DOWSIL™ CY 52-276 Kit), resulting in a substrate stiffness of approximately 2-3 kPa. The gel solution was thoroughly mixed, centrifuged, and degassed before 400 µl were transferred onto a glass coverslip. The solution was spin-coated for 90 s using a homemade spinner, with acceleration and deceleration every 30 seconds to create a smooth, uniform surface. The gels were then cured at 80 °C for 3 h, after which they were stored at RT. To functionalize the silicone surface, the gels were treated with a 7% (v/v) solution of 3-aminopropyl trimethoxysilane (APTES) in pure ethanol for 2 hours. The gels were washed with pure ethanol, a 1:1 ethanol:PBS mixture, and PBS alone, each for three cycles.

As an imaging chamber, an 8-well bottomless sticky-slide (Ibidi, 80828) was mounted on top of each PDMS substrate. Following three washes with PBS (Sigma-Aldrich, D8537), 100 µL of a 100 nm, red-orange fluorescent (540/560) FluoSpheres carboxylate-modified microspheres (Invitrogen, F8800) solution at a dilution of 1:1,000 in PBS was mixed with a final concentration of 100 µg/mL EDC and pipetted onto the top surface of the gel containing 200 µL PBS. The solution was incubated for 20 min at RT and then washed three times with PBS.

To measure traction forces in the under-agarose assay, 300 µl of 1%-agarose was used per well of an 8-well slide with the bottoms coated with the PDMS-TFM substrate described above. At two sites per well, 5×10^3^ *T.gondii-*infected DCs in 1 µl were injected under the agarose layer and allowed to migrate towards a chemokine source of 7 µl 2.5 µg/ml CCL19 in a hole punched with a 3 mm puncher. Live imaging was started 30 min after the injection of DCs and recorded at 30 s per frame.

Image processing was carried out in customized MATLAB scripts. Some crucial functions employed in the workflow are written in *italics* below. First, raw time sequences of fluorescent beads are normalized by *mat2gray* to reduce the brightness differences. We correct the spatial drift resulting from fluctuations of the stage and the microscope by *normxcorr2* for each frame relative to the first frame. Then, aligned image sequences are used for further processing. We select a frame as the reference frame when no cell is in the field of view. We apply *detectMinEigenFeatures* to localize fluorescent beads with subpixel precision at the reference frame. Next, the detected points are fed to a built-in optical flow tracker in MATLAB *vision.PointTracker* to obtain the displacements at each frame relative to the reference. Subsequently, tracking results are structured as input for traction reconstruction. We calculate the traction fields using published software^77^, in which we employ regularized Fourier transform cytometry with optimal estimation of regularization parameters by the Bayesian approach.

#### Atomic force microscopy and Brillouin microscopy

To test the stiffness of the *T. gondii* parasitophorous vacuole, we employed atomic force microscopy (AFM) and Brillouin microscopy (BM). *T. gondii*-infected DCs were let migrate in an under-agarose assay. For this, two parallel holes were punched in the 1%-agarose using a custom-made 2×7 mm puncher at a 2 mm spacing. One hole was loaded with 25 µl of imaging medium supplemented with 0.625 µg/ml CCL19 as chemoattractant, and the other holes were loaded with 25 µl of imaging medium containing 1×10^5^ *T. gondii*-infected DCs stained with only Janelia dye 646 HaloTag ligand (Promega, GA1120). DCs migrated toward the chemokine source, where they squeezed themselves under the 1% agarose gel. After 4 h of migration, the DCs were fixed while still under the agarose layer with 3.7% paraformaldehyde in PBS for 1 h at 37°C, then washed three times with PBS. Then, the agarose was removed to make the cells accessible for AFM and BM, the nuclei were stained with 0.1 μg/ml DAPI (4′,6-Diamidine-2′-phenylindole, #62248, Thermo Scientific), and the dishes were washed three times with PBS.

For AFM measurements, a Nanowizard IV was used (JPK BioAFM, Bruker Nano GmbH) with SNL-10 triangular pyramidal cantilevers (Bruker) with a tip radius of 2-12 nm and a nominal spring constant of 0.35 N/m. Cantilevers were calibrated by the thermal noise method directly inside a glass-bottom petri dish. Quantitative imaging (QI) mode (JPK, Berlin) was used. Cells were scanned with a force load of 3 nN, at 150 µm/s, with a resolution of 2.56 px/µm. The dissipated viscous energy was extracted by calculating the area between the approach and retraction curves in the positive force range for each pixel of the image using a custom-made Python script. Extraction of Young’s modulus and height was performed using the JPKSPM Data Processing software. Here, the Poisson’s ratio was set to 0.5, and the Sneddon model for a triangular pyramidal indenter was applied for the fitting, with a limitation of the indentation curve to 200 nm in the contact region.

Brillouin frequency shift images were obtained, as described before^78^, by employing a Brillouin spectrometer with a two-stage virtually imaged phase array VIPA etalon (Light Machinery, cat. nr. OP-6721-6743-4), a diode laser beam (λ = 780.24 nm, DLC TA PRO 780,

Toptica), a Fabry-Perot interferometer in two-pass configuration and a monochromatic grating (Toptica), a molecular absorption cell filled with rubidium 85Rb (Precision Glassblowing cat nr. TG-ABRB-I85-Q), and a sCMOS camera for the Brillouin spectrum (Teledyne cat nr. Prime BSI). The system was integrated with an inverted microscope stand with motorised stage and focus (AxioObserver 7, Zeiss). For the measurements, a 40x objective lens was used and the exposure time was set to 0.2 s per measurement point. Confocal fluorescence images were acquired in the same region as the Brillouin microscopy with a rescan confocal microscopy module (RCM2, Confocal.nl) with a sCMOS camera (Prime BSI Express, Teledyne), fluorescence excitation with a multi-line laser unit (Skyra, Cobolt, excitation wavelength λ = 405 nm and 637 nm used), and a 50 µm pinhole.

For analysis of the AFM and BM data, epifluorescence (AFM) and confocal microscopy (BM) images were semi-manually aligned with the AFM and BM images, respectively, using a custom-made ImageJ script. Host nuclei and parasitic cargoes were segmented manually based on their respective fluorescence signals, excluding overlapping regions of both the parasite and the host nucleus. Mean BM and AFM values of the host nucleus and parasitic cargo were extracted using a custom-made ImageJ script.

### Image Analysis

#### Software

Fiji (version 2.14.0) ^79^ and LAS X (Version 3.7.6) were used for image processing. Only cells not interacting with other cells, and with single 1-32 stage PVs or uninfected bystanders, were analyzed. For each cell, individual parasite cells inside the PV were counted to assess the PV stage. All data was grouped accordingly. Cell positions were tracked using the ROI manager of Fiji and quantified using custom-made ImageJ scripts. Single-cell data were further analyzed and summarized with R (R version: 4.3.3, R-studio version 2023.12.1.402, tidyverse package version 2.0.0) and exported to GraphPad Prism (version 8.0.1) for statistical analysis.

#### Analysis of migration in straight channels: PV-host nucleus axis, migration speed, PV size

For the orientation of the *Toxoplasma* PV–host nucleus axis, DCs or macrophages were analyzed as they migrated in linear PDMS microchannels of varying widths (8, 10, or 16 µm). The same analysis was performed for DCs carrying beads as well. PV/beads-nucleus orientation and position were documented at the beginning and end of cells migrating ∼500 µm through the microchannels. From this, the frequency of cells with PV-nucleus or bead-nucleus orientations and the frequency of repositioning changes were determined, and the average migration speed over the ∼500 µm distance travelled was calculated for each cell. PV or nucleus size was measured as the length of the longer axis of the PV and the nucleus. Cells were imaged at a constant time interval, depending on the experiment, ranging from 45 to 75 seconds.

#### Analysis of migration in 2 and 4 µm constriction channels

Pore passing dynamics were quantified for DCs or macrophages challenged with *T. gondii,* or for DCs challenged with beads, migrating in linear microchannels of 8 µm width and 5 µm height with a 2 or 4 µm constriction. The passage time was determined by measuring the time between the first time point at which the cell, the nucleus, or the PV entered the constriction on one side and the first time point at which the constriction was fully excited. Further, cells that entered but did not exit the constriction, or that egressed during the 5 h recording period, were documented to calculate the frequency of DC passing, DC being stuck in the constriction, or *T. gondii* egressing. Cells were imaged at 2 min time interval for the fraction of cells passing analysis and at 10 s for the PV-unfolding analysis and analysis of pore passage of the nucleus and PV compared to the whole host cell. For the PV-unfolding analysis, the length of the PV was measured just before the host DC touched the constriction, when the PV touched but did not yet enter the constriction, just after the PV left the constriction, and when the entire host DC left the constriction. Further, the passaging time of single parasites within the PV was measured by determining the time between PV touching both sides of the constriction for the first time, the rear of the first parasite cell passing the middle point of the constriction, and then each subsequent parasite cell passing the middle point of the constriction with its rear.

#### Analysis of EB3-mCherry and Myosin-GFP live fluorescence

DCs derived from EB3-mcherry-or Myosin-GFP-expressing Hoxb8 cells were imaged while migrating through 8 µm-wide, 5 µm-high microchannels. For the microtubule-organizing center (MTOC) positioning analysis, cells with a clearly defined MTOC, as a single brightest spot in the EB3-mCherry fluorescence signal that was consistently present throughout the recording, were analyzed. PV-nucleus-MTOC axis orientation was documented after cells migrated 300-500 µm in the microchannels. The localization of Myosin-GFP was measured using the Fiji line profile over an 8 µm-wide region from the rear to the front of the DC. This was performed by using a custom-made ImageJ script and R script to subtract background signal determined for each replicate, normalize the fluorescence intensity to the average of the entire cell for each cell, resulting in a mean intensity of 1 for each cell, and then the length of each cell was normalized to 0 (rear) to 100 (front).

#### Analysis of blebbing and smooth phenotype

Bone marrow derived *T. gondii*-infected DCs were imaged while migrating in 8 µm wide and 5 µm high microchannels and categorized manually in (1) veiled, for cells with constantly veiled protrusions containing many vesicles, (2) smooth, for cells with a non-veiled cell front not containing vesicles most of the time, or (3) blebbing cells, for cells showing clear signs of frequent quickly developing blebs in some of the frames.

### Computational model

See the extended supplemental file for details.

### Statistics

All experiments were performed at least as three independent biological replicates, and all replicates were validated independently. Statistical analysis was conducted using GraphPad Prism (version 8.0.1) with the appropriate tests for a normal or non-normal distribution, as stated in the figure legend. Fractions of cell subpopulations were calculated from pooled cell numbers of at least three independent biological replicates, and the 95% confidence intervals of the fractions were calculated using the Wilson/Brown hybrid method in Prism. Error bars are defined in the figure legends.

## RESOURCE AVAILABILITY

### Lead contact

Further information and requests for resources and reagents should be directed to and will be fulfilled by the lead contacts, Jörg Renkawitz, Benedikt Sabass, and Javier Periz.

### Materials availability

All newly generated materials associated with the paper are available upon request from the lead contacts.

### Data and code availability

All data supporting the findings of this study are available within the paper. The particle-based model presented in the paper is described in detail in the Supplementary Information and is publicly available: https://github.com/CellMicroMechanics/ToxoDCMigrationSimulationModel. Any additional information required to reanalyze the data reported in this work is available from the lead contacts upon request.

## ACKNOWLEDGEMENTS

We thank Kasia Stefanowski for excellent technical assistance and Janina Kroll for critical reading of the manuscript, Ana-Maria Lennon-Dumenil and Aline Yatim for bone marrow from MyoIIA-Flox*CD11c-Cre mice, Michael Sixt for Myh9-GFP expressing HoxB8 cells, and the Core Facility Bioimaging, the Core Facility Flow Cytometry, and the Animal Core Facility of the Biomedical Center (BMC) for excellent support. This study was supported by the Deutsche Forschungsgemeinschaft (DFG; German Research Foundation; Priority Programme SPP2332, project 492014049, to JR and BS; and DFG project ME 2675/6-2 to JP), the Peter Hans Hofschneider Professorship of the foundation “Stiftung Experimentelle Biomedizin” (to JR), the LMU Institutional Strategy LMU-Excellent within the framework of the German Excellence Initiative (to JR), and the European Research Council (ERC) under the European Union’s Horizon 2020 research and innovation programme (BacForce, G.A.No. 852585, to BS).

## AUTHOR CONTRIBUTIONS

Mauricio J.A. Ruiz-Fernandez: Conceptualization; investigation; methodology; writing – review and editing.

Jianfei Jiang: Software and simulation; methodology; writing – review and editing. Alice Battistella: Investigation; writing – review and editing

Armina Mortazavi: Investigation; methodology; writing – review and editing. Sterre van Wierst: Methodology; writing – review and editing.

Bingzhi Wang: Investigation; writing – review and editing. Artur Kuznetcov: Investigation; writing – review and editing. Rüya Aslan: Investigation; writing – review and editing.

Jack Merrin: Methodology; writing – review and editing.

Daan Vorselen: Supervision; methodology; writing – review and editing. Jochen Guck: Supervision; methodology; writing – review and editing. Markus Meissner: Supervision; methodology; writing – review and editing.

Javier Periz: Conceptualization; supervision; investigation; writing – review and editing.

Benedikt Sabass: Conceptualization; supervision; investigation; funding acquisition; project administration; writing – review and editing.

Jörg Renkawitz: Conceptualization; supervision; investigation; funding acquisition; project administration; writing - original draft; writing – review and editing.

## DECLARATION OF INTEREST

The authors declare no competing interests.

### Declaration of generative AI and AI-assisted technologies in the writing process

The manuscript was edited for grammatical correctness and readability using Grammarly.com and ChatGPT. The AI-assisted modifications were restricted to improving readability, with all scientific content and conclusions remaining the sole responsibility of the authors.

## SUPPLEMENTAL INFORMATION

Supplemental Figures S1 to S5

Supplemental Movies S1 to S11

Supplemental information to numerical model

## SUPPLEMENTAL INFORMATION

### Supplemental Figure Legends

**Supplemental Figure 1. Migration and crossing of microenvironmental barriers by *Toxoplasma*-infected macrophages carrying large parasitic cargoes. (A)** Representative *Toxoplasma gondii*-infected macrophages migrating in linear microchannels are shown as kymographs over time. *Toxoplasma gondii* SAG1-Halo, stained with Janelia dye 646 HaloTag ligand, is shown in red, the host cell shape in yellow, and the macrophage’s nucleus in cyan (Hoechst). Note the different replication stages (1-to 32-stage) and sizes of the parasite within the motile host macrophage. See Figure 1D for quantification of migration speed. **(B)** Quantification of the length of the parasitophorous vacuole in comparison to the host cell nucleus during macrophage migration along linear microchannels, as shown in A). Note that a 4-stage PV equals the length of the host nucleus, while 8-, 16, and 32-staged PVs exceed the size of the macrophage’s nucleus. N = 14 cells (1-stage PV), 19 cells (2-stage PV), 21 cells (4-stage PV), 32 cells (8-stage PV), 15 cells (16-stage PV), and 7 cells (32-stage PV) from 4 independent biological replicates. Paired t-tests. Data are the mean±95% CI. **(C)** Representative dendritic cell carrying a high-stage *Toxoplasma* cargo across a 4-micron-wide pore within a microchannel. *Toxoplasma gondii* is shown in red and the DC nucleus in cyan (Hoechst). See Figure 1F for quantification. **(D)** Passing times of parasitic cargoes of different sizes during macrophage (MC) migration through 2 micrometer-sized pores. Data are the mean±95% CI. Unpaired, non-parametric data: Mann-Whitney test. N = 26 (1-& 2-stage PVs), 34 (4- & 8-stage PVs), 22 (16- & 32-stage PVs) from 4 independent biological replicates.

**Supplemental Figure 2. Intracellular *Toxoplasma* parasites are a major physical bottleneck within motile dendritic cells and macrophages. (A)** Quantification of the frequency of indentation of the dendritic cell (host) nucleus in *Toxoplasma*-infected dendritic cells in the direction of the parasitophorous vacuole (PV), categorized in PVs that are either directly next to the host nucleus, close by (up to 1 micrometer) or distant (more than 1 micrometer). See representative image in Figure 2G. Data are mean±SD, N = 52 cells. **(B)** Live-cell imaging of a representative *Toxoplasma*-infected dendritic cell migrating through a maze-like microenvironment composed of pillars, which connect the lower and upper surfaces and are spaced at 10 micrometer distances. *Toxoplasma gondii* is shown in red, the host cell shape in yellow, and the DC’s nucleus in cyan (Hoechst). Note the dynamic deformations of the host nucleus by the parasitic cargo (see zoom-ins in the yellow boxes). **(C)** Time of passage through 2 micrometer-sized pores of the 1st and 2nd translocating parasite within a parasitophorous vacuole (PV) within a motile dendritic cell. Longer passage times of the 1st parasite in comparison to the 2nd parasite are shown as red connecting lines, whereas the same or shorter passage times are shown in grey and black, respectively. Data are the mean±95%. CI Paired, non-parametric data: Wilcoxon matched-pairs signed rank test. N = 14 (2-stage PVs), 14 (4-stage PVs), 20 (8-stage PVs), 23 (16-stage PVs) from 3 independent biological replicates. **(D)** Representative example of the squeezing of an individual parasite (1-stage parasitophorous vacuole (PV)) within a macrophage during passage through a 2 micrometer-sized pore. *Toxoplasma gondii* is shown in red, the host cell shape in yellow, and the DC’s nucleus in cyan (Hoechst). The yellow dashed square is shown in a zoom. **(E)** Representative 16-stage example showing unfolding of the parasitophorous vacuole (PV) during passage within a macrophage through a 2 micrometer-sized pore. *Toxoplasma gondii* is shown in red, the host cell shape in yellow, and the DC’s nucleus in cyan (Hoechst). The yellow dashed square is shown in a zoom. **(F-H)** Passing times of the host cell body (F), the host nucleus (G), and the parasitophorous vacuole (H) while *Toxoplasma*-infected dendritic cells cross a 2-micrometer pore. Data are the mean±95% CI. Non-paired, non-parametric data: Kruskal-Wallis multiple-comparison test. N = 28 (non-infected bystander DCs), 12 (1-stage PVs), 14 (2-stage PVs), 14 (4-stage PVs), 20 (8-stage PVs), 23 (16-stage PVs) from 3 independent biological replicates. **(I)** Relative passing times of the parasitic cargo and the host nucleus in relation to the host cell passing time. Data are the mean±95% CI. Paired, non-parametric data: Multiple Wilcoxon tests. N = 28 (non-infected bystander DCs), 12 (1-stage PVs), 14 (2-stage PVs), 14 (4-stage PVs), 20 (8-stage PVs), 23 (16-stage PVs) from 3 independent biological replicates.

**Supplemental Figure 3. The intracellular position of the parasitic cargo and its repositioning dynamics change with parasite size. (A)** Representative kymographs of *Toxoplasma*-infected dendritic cells migrating in 10- and 16-micrometer-wide linear channels (1.7 minutes between each frame in the kymograph). *Toxoplasma gondii* is shown in red and the DC’s nucleus in cyan (Hoechst). **(B)** Quantification of the parasitophorous vacuole (PV) to nucleus axis configuration during dendritic cell migration in 10- and 16-micrometer-wide linear channels, as shown in A). N=3 individual biological replicates (20 cells (1-stage PV, 10 µm), 33 cells (2-stage PV, 10 µm), 55 cells (4-stage PV, 10 µm), 38 cells (8-stage PV, 10 µm), 9 cells (16-stage PV, 10 µm) and 11 cells (1-stage PV, 16 µm), 25 cells (2-stage PV, 16 µm), 39 cells (4-stage PV, 16 µm), 49 cells (8-stage PV, 16 µm), 14 cells (16-stage PV, 16 µm)). See also Figure 3B for comparison to 8-micrometer-wide linear channels. **(C)** Positioning of the parasitophorous vacuole (PV) in relation to the host cell nucleus during migration of *Toxoplasma*-infected dendritic cells (DCs) through a maze of pillars (10 micrometer spacing between pillars). See also representative images in Figure S2B. N= 22 cells (1-stage PV), 23 cells (2-stage PV), 17 cells (4-stage PV), 17 cells (8-stage PV), 2 cells (16-stage PV) from 3 independent biological replicates. Data are mean±95% CI. **(D)** Representative kymographs of parasitic cargo repositioning in 10- and 16-micrometer-wide linear channels (3.4 minutes for the 10-micrometer-wide channel, and 1.7 minutes for the 16-micrometer-wide channel between each frame in the kymograph). *Toxoplasma gondii* is shown in red and the DC’s nucleus in cyan (Hoechst). **(E)** Quantification of the dynamic repositioning of PV cargoes towards the host cell front, frontward of the host nucleus, as shown in D). N=3 individual biological replicates (20 cells (1-stage PV, 10 µm), 33 cells (2-stage PV, 10 µm), 56 cells (4-stage PV, 10 µm), 39 cells (8-stage PV, 10 µm), 9 cells (16-stage PV, 10 µm), and 12 cells (1-stage PV, 16 µm), 27 cells (2-stage PV, 16 µm), 40 cells (4-stage PV, 16 µm), 50 cells (8-stage PV, 16 µm), 14 cells (16-stage PV, 16 µm)).

**Supplemental Figure 4. Immune cell motility with non-deformable and deformable bead cargoes. (A)** Representative dendritic cell (DC; nucleus in cyan) carrying a 2-stage parasitic cargo (red) and a 6 micron-sized rigid bead (green) migrating through the maze of a 3D pillar forest. **(B)** Quantification of the PV to nucleus axis configuration in relation to rigid latex beads, as shown in A). N=21 cells from 3 individual biological replicates. Data are shown as violin plots with the median. **(C)** Representative dendritic cell (DC) with a cargo composed of two deformable (6.5 kPa) beads, migrating in a linear microchannel with a two-micrometer pore. Deformable beads in green and dendritic cell nucleus in cyan. Note the deformation of the DC nucleus by the deformable bead. **(D)** Transport and positioning of deformable beads (as in C) within motile DCs along linear microchannels. These representative cells show examples of forward and rearward positioning of the beads relative to the host nucleus within cells carrying small (1 bead), intermediate (7 beads), or large (11 beads) cargoes of deformable beads. See Figure 3F for quantification. **(E)** Representative example of EB3-mcherry expressing dendritic cells (white; label for the MTOC) infected with *Toxoplasma gondii* shown in red. The host nucleus is shown in cyan, and the position of the MTOC is marked with a purple arrow. **(F)** Quantification of the intracellular configuration of the DC MTOC, DC nucleus, and parasite axis, showing that most cells position the parasite together with the host nucleus forward of the host MTOC. N=3 individual biological replicates (19 cells (1-stage PV, 10 µm), 26 cells (2-stage PV, 10 µm), 22 cells (4-stage PV, 10 µm), 44 cells (8-stage PV, 10 µm), 18 cells (16-stage PV, 10 µm), and 5 cells (32-stage PV, 16 µm).

**Supplemental Figure 5. Adaptation of Host Cell Forces to the Size of the Parasitic Cargo. (A)** Representative examples of Lifeact-GFP-expressing dendritic cells carrying different stages of *Toxoplasma gondii* (small 2-stage and large 8-stage cargoes, as well as uninfected bystander cell). Zoom-ins in the yellow boxes highlight the different distributions of actin within the protrusions. Note the reduced actin signal within bleb-like protrusions in the dendritic cells carrying a large 8-stage parasite. **(B)** Representative images of alpha-tubulin immunofluorescence stainings (white) of control (DMSO) or microtubule-inhibited (nocodazole) dendritic cells (nucleus in blue; DAPI) infected with *Toxoplasma gondii* (red; SAG1-Halo). **(C)** Quantification of the blebbing and smooth cell front phenotype in the presence of the microtubule inhibitor nocodazole or DMSO controls. N=5 individual biological replicates (16 cells (1-stage PV, DMSO), 17 cells (2-stage PV, DMSO), 13 cells (4-stage PV, DMSO), 33 cells (8-stage PV, DMSO), 42 cells (16-stage PV, DMSO), and 4 cells (32-stage PV, DMSO), and 13 cells (1-stage PV, nocodazole), 11 cells (2-stage PV, nocodazole), 13 cells (4-stage PV, nocodazole), 26 cells (8-stage PV, nocodazole), 31 cells (16-stage PV, nocodazole), and 3 cells (32-stage PV, nocodazole)). **(D)** Representative image (left) and heat map (right) of TFM-beads and their displacement during the migration of *Toxoplasma*-infected dendritic cells. **(E)** Quantification of D, comparing beads displacements in wild-type (WT) and myosin-IIA knockout dendritic cells. N = 29 cells (1-stage PV, Control), 23 cells (2-stage PV, Control), 23 cells (4-stage PV, Control), 29 cells (8-stage PV, Control), 36 cells (16-stage PV, Control), and 14 cells (32-stage PV, Control), and 6 cells (1-stage PV, Myosin-IIA KO), 5 cells (2-stage PV, Myosin-IIA KO), 8 cells (4-stage PV, Myosin-IIA KO), 19 cells (8-stage PV, Myosin-IIA KO), 22 cells (16-stage PV, Myosin-IIA KO), and 10 cells (32-stage PV, Myosin-IIA KO) from 5 independent biological replicates. Data are the mean±SEM.

### Supplemental Movies

**Supplemental Movie S1. Transport of *Toxoplasma*-cargoes by infected dendritic cells through confining microenvironments.** Representative dendritic cells (DCs) infected with *Toxoplasma gondii* were imaged while migrating along confining, linear microchannels (8 micron wide and 5 microns high) towards a CCL19 chemokine gradient. Note the different replication stages (1, 2, 4, 8, 16, and 32-stages) and thus sizes of the parasitic cargo and that the parasitic cargoes can position in front and in the rear of the host. *Toxoplasma gondii* parasites expressing SAG1-Halo are shown in red (Janelia dye 646 Halo ligand) and the DC nuclei are shown in cyan (Hoechst 33342). Time is indicated as minutes (min).

**Supplemental Movie S2. Transport of *Toxoplasma*-cargoes by infected macrophages through confining microenvironments.** Representative macrophages infected with *Toxoplasma gondii* were imaged while migrating along confining, linear microchannels (8 micron wide and 5 microns high) towards a CCL19 chemokine gradient. Note the different replication stages (1, 2, 4, 8, 16, and 32-stages) and thus sizes of the parasitic cargo. *Toxoplasma gondii* parasites expressing SAG1-Halo are shown in red (Janelia dye 646 Halo ligand) and the macrophages nuclei are shown in cyan (Hoechst 33342). Time is indicated as minutes (min).

**Supplemental Movie S3. *Toxoplasma*-cargoes passing 2 micrometer-wide pores during their transport in infected dendritic cells.** Representative dendritic cells (DCs) infected with *Toxoplasma gondii* were imaged while migrating through a two micrometer-wide pore within confining, linear microchannels (8 micron wide and 5 microns high) towards a CCL19 chemokine gradient. Note the different replication stages (1, 2, 4, 8, and 16-stages) and thus sizes of the parasitic cargo. Individual parasites squeeze and deform during the crossing of this microenvironmental barrier. The representative 2- to 16-stage parasitic cargoes show unfolding and refolding during the pore passage, allowing individual parasites to pass one-by-one. *Toxoplasma gondii* parasites expressing SAG1-Halo are shown in red (Janelia dye 646 Halo ligand) and the DC nuclei are shown in cyan (Hoechst 33342). Time is indicated as minutes (min).

**Supplemental Movie S4. *Toxoplasma*-cargoes passing 2 micrometer-wide pores during their transport in infected macrophages.** Representative macrophages infected with *Toxoplasma gondii* were imaged while migrating through a two micrometer-wide pore within confining, linear microchannels (8 micron wide and 5 microns high) towards a CCL19 chemokine gradient. Note the different replication stages (1, 2, 4, 8, and 16-stages) and thus sizes of the parasitic cargo. Individual parasites squeeze and deform during the crossing of this microenvironmental barrier. The representative 2- to 16-stage parasitic cargoes show unfolding and refolding during the pore passage, allowing individual parasites to pass one-by-one. *Toxoplasma gondii* parasites expressing SAG1-Halo are shown in red (Janelia dye 646 Halo ligand) and the macrophage nuclei are shown in cyan (Hoechst 33342). Time is indicated as minutes (min).

**Supplemental Movie S5. Active repositioning of *Toxoplasma*-cargoes towards the cell fronts of infected, motile dendritic cells.** Representative dendritic cells (DCs) infected with *Toxoplasma gondii* were imaged while migrating along confining, linear microchannels (8 micron wide and 5 microns high) towards a CCL19 chemokine gradient. Note that the parasitic cargoes can reposition from the cell front to behind the host nucleus (1-stage) or overtake the host nucleus to reposition towards the cell front (4- to 32-stage). *Toxoplasma gondii* parasites expressing SAG1-Halo are shown in red (Janelia dye 646 Halo ligand) and the DC nuclei are shown in cyan (NucBlue, Hoechst 33342). Time is indicated as minutes (min).

**Supplemental Movie S6. Migration of dendritic cells carrying deformable bead cargoes.** Representative dendritic cells (DCs) migrating along confining, linear microchannels (8 micron wide and 5 microns high) while carrying different numbers of deformable beads (1, 2, 4, 7, and 11 beads; green). The DC nucleus (Hoechst) is shown in blue. The representative examples show the two possible configurations of beads and the DC nucleus along the cell axis (beads in front or rear of the DC nucleus). Time is indicated as minutes (min).

**Supplemental Movie S7. Positioning of the parasitic cargo relative to the microtubule-organising center.** Representative dendritic cells (DCs) expressing EB3-mCherry, as a microtubule-organising center (MTOC) marker, infected with *Toxoplasma gondii* were imaged while migrating along confining, linear microchannels (8 micron wide and 5 microns high) towards a CCL19 chemokine gradient. Representative uninfected bystander DCs and DCs carrying 1- or 4-stage parasitic cargoes are shown in the most prominent MTOC-nucleus (uninfected) or parasite-MTOC-nucleus conformations. Lastly, a DCs carrying a 16-stage parasitic cargo is shown, demonstrating how the parasite can overtake the host MTOC while still behind the host nucleus, notably also incurring some deformation of the parasitic cargo for it to fit next to the MTOC. EB3 microtubules tips and MTOC visible in grey (bottom videos, EB3-mCherry), *Toxoplasma gondii* parasites expressing SAG1-Halo are shown in red (Janelia dye 646 Halo ligand) and the DC nuclei are shown in cyan (NucBlue, Hoechst 33342). The positioning of the parasite to the host MTOC and nucleus is indicated at the top and the time is indicated in minutes (min).

**Supplemental Movie S8. Simulating the intracellular localization of the parasitic cargo and the host nucleus.** Representative movie from the computational simulation, showing the parasitic cargo in red, the host cytoplasm in gray, the host cell cortex in orange and the host nucleus in blue (first movie part). The second movie part shows additionally the intracellular fluid particle density. See main text, main figures, and the Supplemental Information for further details.

**Supplemental Movie S9. Localization of myosin during the migration of dendritic cells carrying parasitic or deformable bead cargoes.** Representative myosin-GFP (white) expressing dendritic cells (DC) migrating along confining, linear microchannels while carrying different parasitic stages (red; 1- and 8-staged) or different numbers of deformable beads (green; 1 and 9 beads). The DC nucleus (Hoechst) is shown in blue. Note the rearward localized myosin accumulation in the DC carrying a large, 8-staged parasitic cargo. Time is indicated as minutes (min).

**Supplemental Movie S10. Blebbing cell protrusions in dendritic cells carrying large parasitic cargoes.** First part: Representative dendritic cells (DC) migrating along confining, linear microchannels while carrying different sizes of parasitic cargo (red). Note different protrusion shapes, including bleb-like protrusions. Second part: Representative Lifeact-GFP (green) expressing dendritic cell (DC). migrating along confining, linear microchannels while carrying a large parasitic cargo (red). Note the quick extension of the bleb that is initially devoid of a Lifeact-GFP signal. Time is indicated as minutes (min).

**Supplemental Movie S11. Forces onto the environment by *Toxoplasma gondii*-infected dendritic cells.** Representative example of traction force microscopy, showing dendritic cells (DC) carrying either a small, 2-stage or a large, 16-stage parasitic cargo. Traction vectors are shown in orange, *Toxoplasma gondii* parasites expressing SAG1-Halo are shown in red (Janelia dye 646 Halo ligand), and the DC nuclei are shown in cyan (NucBlue, Hoechst 33342). Time is indicated as minutes (min).

Supplemental Movie S12. Impaired translocation of *Toxoplasma*-cargoes through narrow pores upon myosin inhibition, and when carrying deformable beads. First part: Representative dendritic cell (DC) infected with *Toxoplasma gondii*, and migrating through a 4 micrometer-sized pore in the presence of the myosin inhibitor para-nitroblebbistatin. Second part: Representative dendritic cell (DC) infected with *Toxoplasma gondii*, trying to migrate through a 2 micrometer-sized pore in the presence of the myosin inhibitor para-nitroblebbistatin. Third part: Representative dendritic cell (DC) carrying deformable beads as a cargo, and trying to migrate through a 2 micrometer-sized pore. Fourth part: Representative parasitic egress at a 2 micrometer-sized pore. The movies show representative cells from at least three independent biological replicates. Time is indicated as minutes (min).

## Notes

### Competing Interest Statement

The authors have declared no competing interest.

### Summary of Updates

The revised manuscript (i) extends the findings to different immune cell types using distinct migratory strategies, (ii) performs multiple, complementary physical measurements of cargoes, (iii) computationally simulates the identified reorganisation in cellular architecture and dynamics, and (iv) systematically compares parasitic cargoes with parasite-independent foreign cargoes.

