## Supplemental Figure 1 for "Size-Dependent Mechanoadaptation Enables Migration with Oversized Parasitic Cargo"

### Figure S1. Migration and Crossing of Microenvironmental Barriers by *Toxoplasma*-infected Macrophages Carrying Large Parasitic Cargoes.

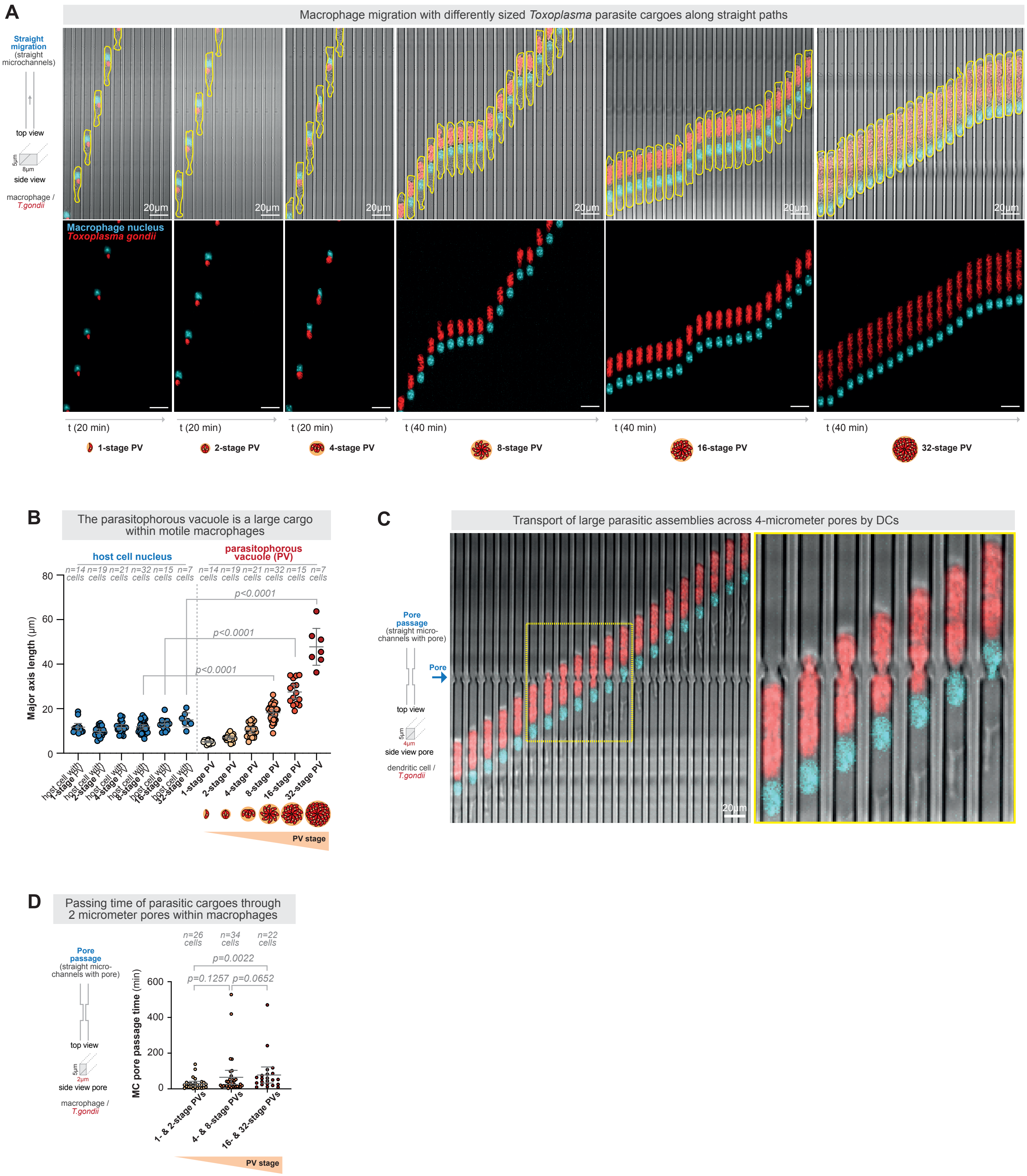
