## Supplemental Figure 2 for "Size-Dependent Mechanoadaptation Enables Migration with Oversized Parasitic Cargo"

### Figure S2. Intracellular *Toxoplasma* Parasites are a Major Physical Bottleneck within Motile Dendritic Cells and Macrophages.

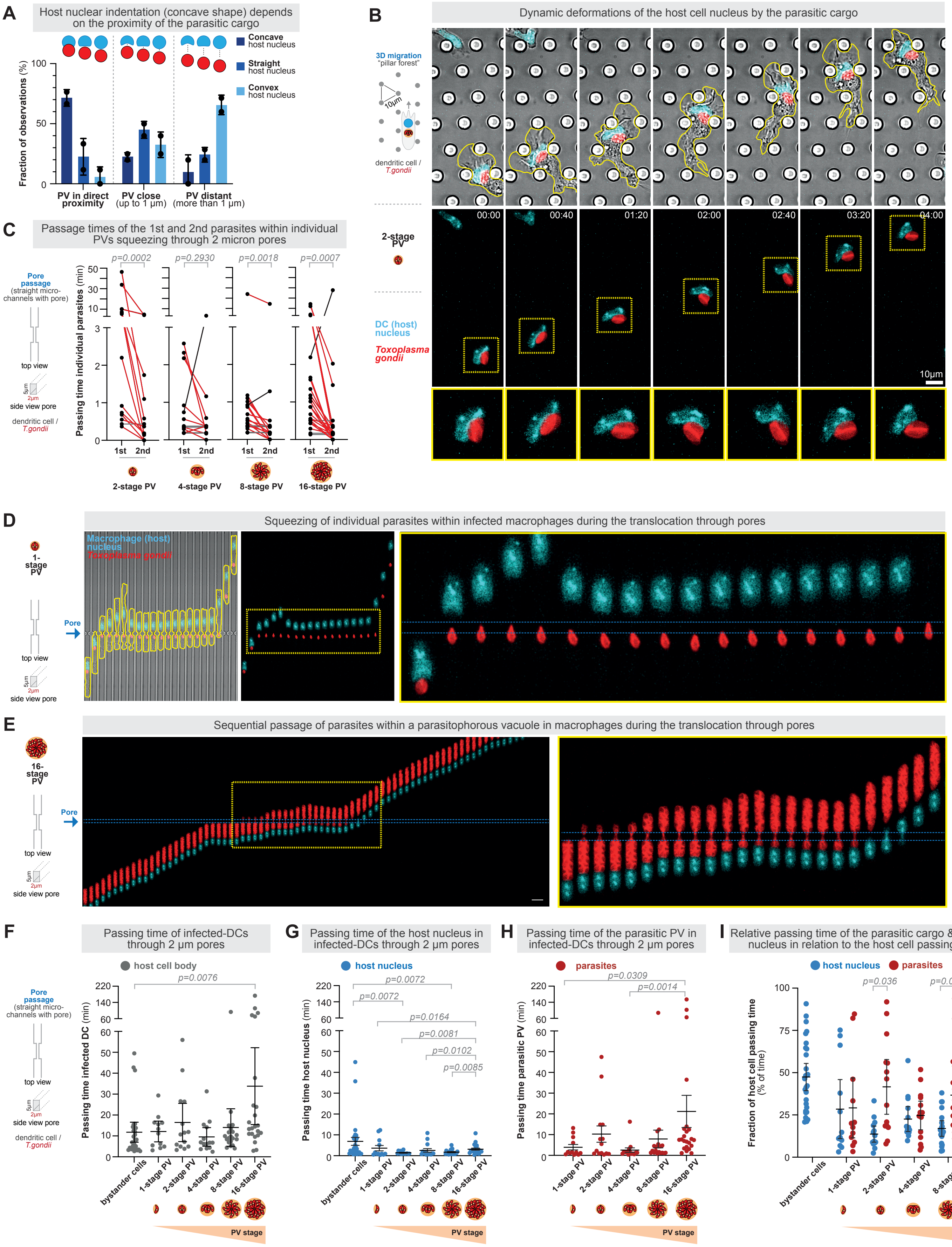
