## Supplemental Figure 3 for "Size-Dependent Mechanoadaptation Enables Migration with Oversized Parasitic Cargo"

**Figure S3. The Intracellular Position of the Parasitic Cargo and its Repositioning Dynamics Change with Parasitic Size.**

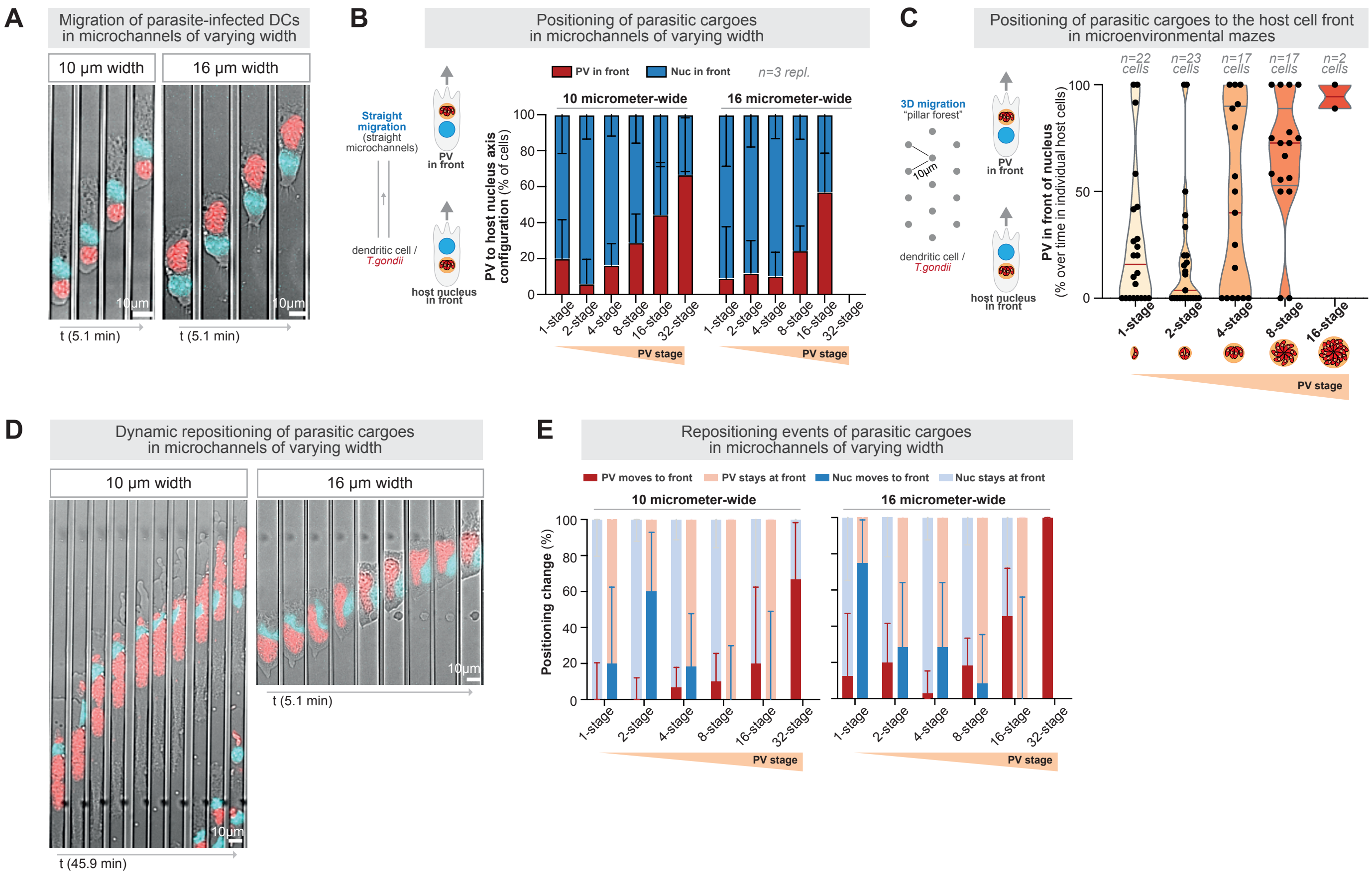
