## Supplementary figures and images for "Size-Dependent Mechanoadaptation Enables Migration with Oversized Parasitic Cargo"

### Supplemental Figure 4

#### Figure S4. Immune cell motility with non-deformable and deformable bead cargoes

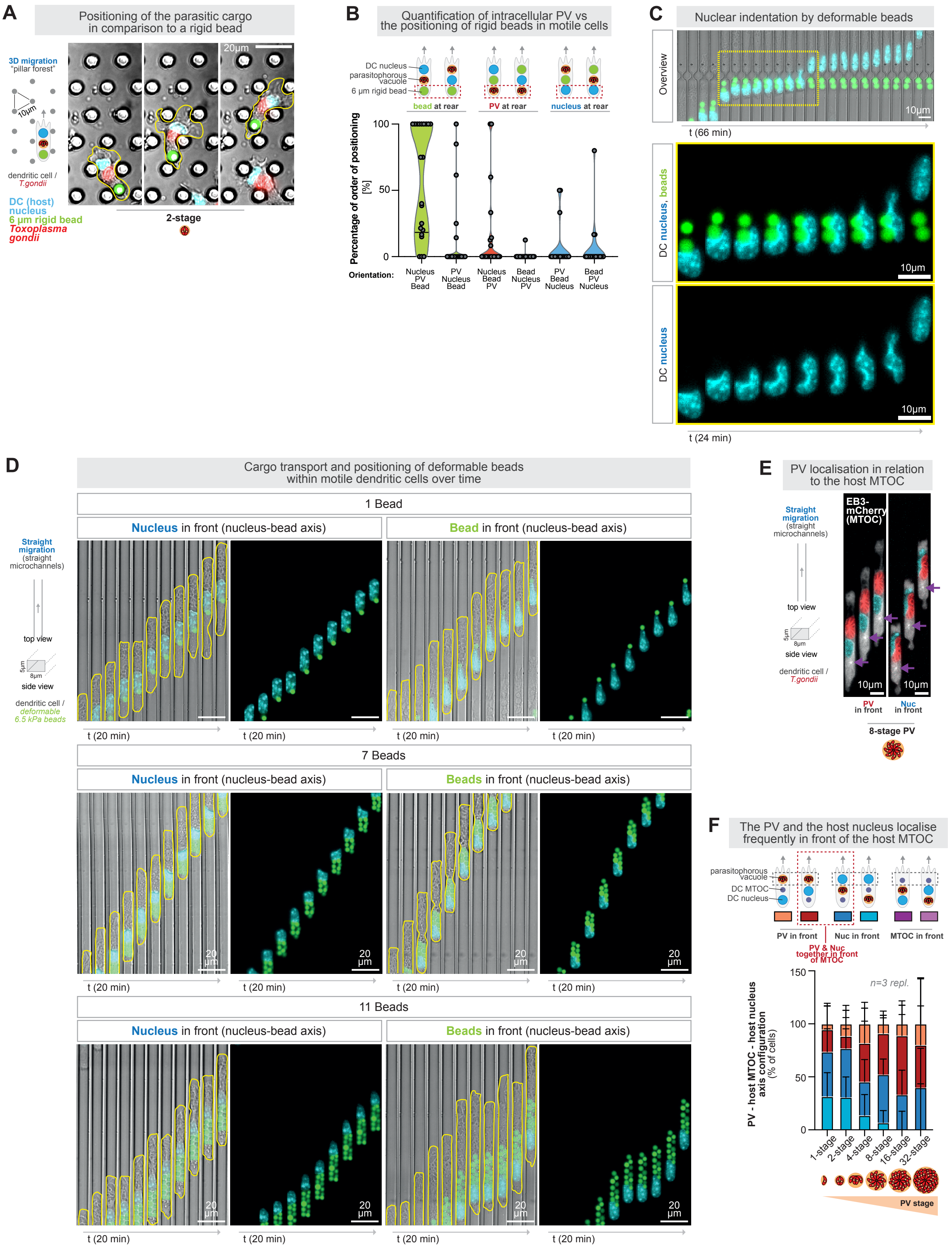

### Supplemental Figure 5

#### Figure S5. Adaptation of Host Cell Forces to the Size of the Parasitic Cargo.

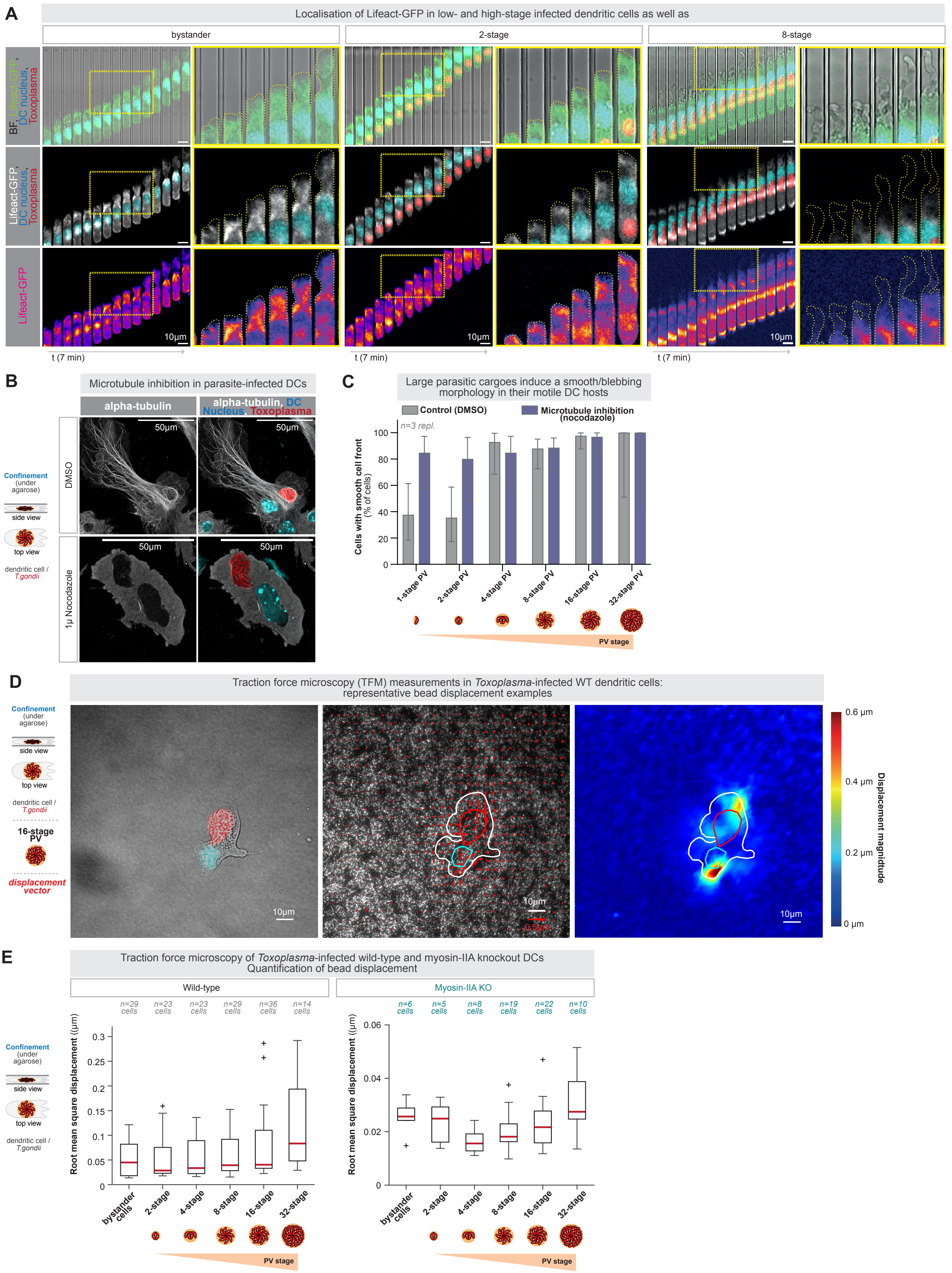
