## Supplemental Information to the Computational Model for "Size-Dependent Mechanoadaptation Enables Migration with Oversized Parasitic Cargo"

### Supplementary Information for Size-Dependent Mechanoadaptation Enables Migration with Oversized Parasitic Cargo

This supplementary information provides a detailed description of the particle-based model presented in the main manuscript, along with additional results and discussions that expand the main findings.

#### 1 The model

We employ a two-dimensional geometry, in which the outer cell boundaries are composed of spring-connected particles with enclosed fluid particles. We simulate the ensemble of cell plasma membrane and f-actin cortex in a coarse-grained way with a single layer of membrane particles that interact with the external environment via local friction. To study the infected host-cell system, we represent the intracellular organelles, i.e., nucleus and the parasitophorous vacuole (PV), as elastic objects [1, 2], and employ dissipative particle dynamics (DPD) [3, 4, 5] to simulate the cytoplasmic fluid. No-flux boundary conditions are imposed across object boundaries by proper spacing between particles.

##### 1.1 Dissipative particle dynamics

DPD is a method for a particle-based, mesoscopic description of hydrodynamic systems. We utilize this coarse-grained fluid to mimic intracellular flow driven by cortical dynamics. Fluid particles experience pairwise interactions with their neighbors within a cutoff radius $r_c$. For a pair of particles $i$ and $j$, we define the relative position as $\mathbf{r}_{ij} = \mathbf{r}_i - \mathbf{r}_j$, its magnitude $r_{ij} = |\mathbf{r}_{ij}|$, and the corresponding unit vector $\hat{\mathbf{r}}_{ij} = \mathbf{r}_{ij}/r_{ij}$. The relative velocity between two particles is $\mathbf{v}_{ij} = \mathbf{v}_i - \mathbf{v}_j$. The force on particle $i$ due to particle $j$ consists of a conservative force $\mathbf{F}_{C,ij}$, a dissipative force $\mathbf{F}_{D,ij}$, and a random force $\mathbf{F}_{R,ij}$. These forces are calculated as

$$\mathbf{F}_{C,ij} = a\omega(r_{ij})\hat{\mathbf{r}}_{ij}, \tag{1}$$

$$\mathbf{F}_{D,ij} = -\gamma\omega^2(r_{ij})(\hat{\mathbf{r}}_{ij} \cdot \mathbf{v}_{ij})\hat{\mathbf{r}}_{ij}, \tag{2}$$

$$\mathbf{F}_{R,ij} = \sigma\omega(r_{ij})\zeta_{ij}(\Delta t)^{-1/2}\hat{\mathbf{r}}_{ij}, \tag{3}$$

where the constant $a$ determines the strength of the conservative repulsive force that controls the compressibility of the fluid, $\omega(r_{ij})$ is a weighting function given by $\omega(r_{ij}) = 1 - \frac{r_{ij}}{r_c}$ for $r_{ij} < r_c$ and zero otherwise, $\hat{\mathbf{r}}_{ij}$ is a unit vector in the direction of $\mathbf{r}_i - \mathbf{r}_j$, $\gamma$ is the friction coefficient, $\Delta t$ is the time step, and $\zeta_{ij}$ is a Gaussian random number with zero mean and unit variance. The parameter $\sigma$, which fixes the magnitude of the noise, is set to $\sqrt{2k_BT\gamma}$, ensuring that the fluctuation-dissipation condition is satisfied in the fluid.

#### 1.2 Interface dynamics

We treat the cortex and the organelles as two-dimensional elastic objects whose shape evolution is governed by a shape energy that is frequently used in vertex models [2, 6]. The shape energy is written as

$$E_{\text{Shape}} = E_A + E_P + E_{\text{bending}} + E_{\text{spring}} + E_{\text{tension}}, \tag{4}$$

$$E_A = \frac{1}{2}K_A(A - A_0)^2, \tag{5}$$

$$E_P = \frac{1}{2}K_P(P - P_0)^2, \tag{6}$$

$$E_{\text{bending}} = \frac{1}{2}\sum_{i=1}^{n} K_C(\theta_i - \theta_0)^2, \tag{7}$$

$$E_{\text{spring}} = \frac{1}{2}\sum_{i=1}^{n} K_l(l_i - l_0)^2, \tag{8}$$

$$E_{\text{tension}} = \sum_{i=1}^{n} \alpha_i l_i, \tag{9}$$

where $A$,$P$,$\theta_i$,$l_i$ are functions of relevant particle positions. Equations (5) and (6) represent the energy penalties associated with deviations of the area $A$ enclosed by the interface and perimeter $P$ of the interface from their rest values $A_0$ and $P_0$. The strength of these penalties is determined by elastic constants $K_A$ and $K_P$. Equations (7) and (8) account for the local bending rigidity and stretchability of the interface, with elasticity constants $K_C$ and $K_l$, penalizing deviations from a rest angle $\theta_0$ and the length of the edge $l_0$. Additionally, Eq. (9) models the line tension along the cortex, where a graded tension coefficient $\alpha_i$ induces localized contraction or expansion, allowing contractility. Note that Eq. (9) is only used to model the cortex, while Eqs. (5) - (8) are sufficient for the description of the nucleus and the PV. The net force on the $i$-th particle, resulting from the tendency to reduce the shape energy, is given by the negative gradient as

$$\mathbf{F}_{\text{Shape},i} = -\nabla_{\mathbf{r}_i} E_{\text{Shape}}. \tag{10}$$

Considering that the computational cost of the calculation of the numerical energy gradients at each time step is high, we derive analytic energy gradients below to reduce the simulation time. The area enclosed by the interface, consisting of a sequence of $n$ particles, is given by

$$A = \frac{1}{2}\sum_{i}(x_i y_{i+1} - x_{i+1} y_i), \tag{11}$$

where index $i$ increases with counterclockwise ordered particles and we set $i = 1$ when $i = n + 1$ since the polygon is closed. Then the gradients of the area energy at $(x_i, y_i)$ read

$$\frac{\partial E_A}{\partial x_i} = \frac{1}{2} K_A (A - A_0)(y_{i+1} - y_{i-1}), \tag{12}$$

$$\frac{\partial E_A}{\partial y_i} = \frac{1}{2} K_A (A - A_0)(x_{i-1} - x_{i+1}). \tag{13}$$

The perimeter of the interface is written as $P = \sum_i^n l_i$ with local edge length

$$l_i = \sqrt{(x_i - x_{i+1})^2 + (y_i - y_{i+1})^2}. \tag{14}$$

Applying the chain rule to Eq. (6) with respect to $(x_i, y_i)$ gives

$$\begin{aligned} \frac{\partial E_P}{\partial x_i} &= K_P (P - P_0) \frac{\partial P}{\partial x_i} \\ &= K_P (P - P_0) \left( \frac{x_i - x_{i+1}}{l_i} - \frac{x_i - x_{i-1}}{l_{i-1}} \right), \end{aligned} \tag{15}$$

$$\begin{aligned} \frac{\partial E_P}{\partial y_i} &= K_P (P - P_0) \frac{\partial P}{\partial y_i} \\ &= K_P (P - P_0) \left( \frac{y_i - y_{i+1}}{l_i} - \frac{y_i - y_{i-1}}{l_{i-1}} \right), \end{aligned} \tag{16}$$

where the global nature of $E_P$ enters only through the scalar factor $(P - P_0)$, while the geometric derivative $\partial P / \partial (x_i, y_i)$ is local since $(x_i, y_i)$ appears only in the two adjacent edges $l_i$ and $l_{i-1}$. While Eqs. (15) and (16) determine the restoration force on particles due to the perimeter change, the spring energy, Eq. (8), prevents local distortions on the edge $\mathbf{l}_i = \big(x_{i-1} - x_i, y_{i-1} - y_i\big)$ with the length of $l_i = \sqrt{(x_{i-1} - x_i)^2 + (y_{i-1} - y_i)^2}$ from its rest length $l_0$. The x-component of the energy gradient is calculated as

$$\begin{aligned} \frac{\partial E_{\text{spring}}}{\partial x_i} &= K_l (l_i - l_0) \frac{\partial l_i}{\partial x_i} + K_l (l_{i-1} - l_0) \frac{\partial l_{i-1}}{\partial x_i} \\ &= K_l \Bigg( (1 - \frac{l_0}{l_i})(x_i - x_{i+1}) + (1 - \frac{l_0}{l_{i-1}})(x_i - x_{i-1}) \Bigg), \end{aligned} \tag{17}$$

$$\tag{18}$$

and similar for the y-component

$$\frac{\partial E_{\text{spring}}}{\partial y_i} = K_l \Bigg( (1 - \frac{l_0}{l_i})(y_i - y_{i+1}) + (1 - \frac{l_0}{l_{i-1}})(y_i - y_{i-1}) \Bigg). \tag{19}$$

To express the bending energy gradient, we define the angle $\theta_i$ between two adjacent edges $\mathbf{l}_{i-1}$ and $\mathbf{l}_i$ by

$$\cos \theta_i = \frac{\mathbf{l}_{i-1} \cdot \mathbf{l}_i}{l_{i-1} l_i}, \tag{20}$$

$$\theta_i = \arccos \frac{\mathbf{l}_{i-1} \cdot \mathbf{l}_i}{l_{i-1} l_i}, \tag{21}$$

with its derivative with respect to $x_i$

$$\begin{aligned} \frac{\partial \theta_i}{\partial x_i} &= - \frac{1}{\sqrt{1 - \cos^2 \theta_i}} \frac{\partial \cos \theta_i}{\partial x_i} \\ &= - \frac{1}{\sqrt{1 - \cos^2 \theta_i}} \frac{\partial}{\partial x_i} \left( \frac{\mathbf{l}_{i-1} \cdot \mathbf{l}_i}{l_{i-1} l_i} \right). \end{aligned} \tag{22}$$

Substitution of the expressions for $\mathbf{l}_{i-1}$, $\mathbf{l}_i$, $l_{i-1}$ and $l_i$ into Eq. (22) yields after some algebra

$$\frac{\partial E_{\text{bending}}}{\partial x_i} = -\frac{K_C(\theta_i-\theta_0)}{\sqrt{1-\cos^2\theta_i}}\Big( (\frac{\cos\theta_i}{l_{i-1}^2}-\frac{1}{l_{i-1}l_i})(x_i-x_{i-1})+(\frac{\cos\theta_i}{l_i^2}-\frac{1}{l_{i-1}l_i})(x_i-x_{i+1})\Big), \quad (23)$$

and similarly for the y-component

$$\frac{\partial E_{\text{bending}}}{\partial y_i} = -\frac{K_C(\theta_i-\theta_0)}{\sqrt{1-\cos^2\theta_i}}\Big( (\frac{\cos\theta_i}{l_{i-1}^2}-\frac{1}{l_{i-1}l_i})(y_i-y_{i-1})+(\frac{\cos\theta_i}{l_i^2}-\frac{1}{l_{i-1}l_i})(y_i-y_{i+1})\Big). \quad (24)$$

The above expressions are adopted in the representation of all interface-forming objects in our simulations, including the nucleus, the PV, and the cortex. In addition, we consider an active tension and the resulting flow in the cortex. For the tension energy, we assume that the dynamics setting up the cortical tension is much faster than the dynamics of the cell migration, and the coefficients $\alpha_i$ that determine the active tension are assumed to be constant. By taking derivatives, we obtain

$$\frac{\partial E_{\text{tension}}}{\partial x_i} = \frac{\alpha_{i-1}}{l_{i-1}}(x_i-x_{i-1})+\frac{\alpha_i}{l_i}(x_i-x_{i+1}), \quad (25)$$

$$\frac{\partial E_{\text{tension}}}{\partial y_i} = \frac{\alpha_{i-1}}{l_{i-1}}(y_i-y_{i-1})+\frac{\alpha_i}{l_i}(y_i-y_{i+1}). \quad (26)$$

To account for the cell contractility resulting from inhomogeneous myosin distribution, we employ a spatially varying profile of the line tension coefficient $\alpha_i$ given by

$$\alpha_i = \tau_0 + \tau_\alpha(1-\cos\phi_i), \quad (27)$$

where $\tau_0$ and $\tau_\alpha$ scale the cortical tension, $\phi_i$ is the polar angle between the position vector of a cortical particle $i$ and the predefined cell polarity in the cortex center-of-mass frame [6]. For simplicity, we set the cell to a constant polarity along the $x$ direction. This cortical tension profile leads to a high tension region at the cell rear and a low tension region at the cell front, modelling the actomyosin contraction in the rear and actin polymerization at the leading edge, respectively. Additionally, the graded cortical tension contributes to the cortical flow that drives the cell migration [7, 8].

To drive a stable cortical flow, the exchange of matter through actin polymerization and depolymerization must be considered. In our model, polymerization and depolymerization are represented as the addition to, or removal of, a single cortical particle. Both processes share the same global rate $\kappa$, so that, on average, the total number of cortex particles remains constant. During each time step of length $\Delta t$, we perform two independent Bernoulli trials, one for polymerization and one for depolymerization, each with success probability $P = 1-\exp(-\kappa\,\Delta t)$. Because we choose a small time step such that $\kappa\Delta t \ll 1$, we have $P \simeq \kappa\Delta t$ and the probability of two or more occurrences is $\mathcal{O}\big((\kappa\Delta t)^2\big)$-an order of magnitude smaller. We therefore interpret the success as exactly one event of polymerization or depolymerization. If a polymerization trial succeeds, a new particle is inserted at the midpoint of the longest edge, whereas a successful depolymerization trial removes the particle associated with the shortest edge.

##### 1.3 Additional interactions

In addition to the forces governing interface dynamics and intracellular fluid, mechanical coupling between the nucleus, cortex, and external environment is necessary to correctly capture cell migration dynamics. It is known that the nucleus is mechanically tethered by the linker of the nucleoskeleton and cytoskeleton (LINC) complex [9], which plays a role in the nuclear positioning of cortical-flow-driven migratory cells [10]. We focus on two net effects of this coupling. First, the nucleus tends to stay at an equilibrium position. Second, the nucleus can be displaced by the retrograde cortical flow. To account for the first effect, a spring-like force is employed between the center of mass of the nucleus and the cortex. To map with our particle representation, the force is uniformly redistributed to particles on the nuclear membrane and the cortex, which reads

$$\mathbf{F}_{\text{nucleus},i} = -\frac{K}{N_{\text{nucleus}}}(\mathbf{r}_{c,\text{nucleus}} - \mathbf{r}_{c,\text{cortex}}), \tag{28}$$

$$\mathbf{F}_{\text{cortex},j} = \frac{K}{N_{\text{cortex}}}(\mathbf{r}_{c,\text{nucleus}} - \mathbf{r}_{c,\text{cortex}}), \tag{29}$$

where $K$ is the coupling strength, $N$ denotes the number of particles in the related membrane, and the $\mathbf{r}_c$ are the centers of mass of the interface-forming structures. Equation (29) provides the equal-and-opposite force required by Newton's third law. Furthermore, the coupling between the nucleus and retrograde flow is modelled by a friction-like force, which slows down the relative motion between the flowing cortical particles and nuclear particles. We use the following forms

$$\mathbf{f}_{\text{nucleus},i} = -\xi_{\|}\big(\mathbf{v}_{ij} \cdot \hat{\mathbf{r}}_{ij}\big)\hat{\mathbf{r}}_{ij} - \xi_{\perp}(\mathbf{I} - \hat{\mathbf{r}}_{ij} \otimes \hat{\mathbf{r}}_{ij})\mathbf{v}_{ij}, \tag{30}$$

$$\mathbf{f}_{\text{cortex},j} = -\mathbf{F}_{\text{nucleus},i}. \tag{31}$$

Here $\hat{\mathbf{r}}_{ij}$ is the unit vector from a cortical particle $j$ to a nuclear particle $i$ and $\mathbf{v}_{ij}$ is the relative velocity. The first term in Eq. (30) damps the relative motion parallel to $\hat{\mathbf{r}}_{ij}$ with coefficient $\xi_{\|}$. The projector $\mathbf{I} - \hat{\mathbf{r}}_{ij} \otimes \hat{\mathbf{r}}_{ij}$ extracts the tangential component of the relative motion, where damping is determined by the coefficient $\xi_{\perp}$. Equation (31), again, accounts for Newton's third law. The total coupling force between the nucleus and the cortex, therefore, is given by

$$\mathbf{F}_{\text{coupling},i} = \begin{cases} \mathbf{F}_{\text{nucleus},i} + \mathbf{f}_{\text{nucleus},i}, & i \in \text{nucleus}, \\ \mathbf{F}_{\text{cortex},i} + \mathbf{f}_{\text{cortex},i}, & i \in \text{cortex}. \end{cases} \tag{32}$$

The interaction between the cortex and the environment is formulated similarly to

$$\mathbf{F}_{\text{wall},i} = -\eta_{\|}(\mathbf{I} - \mathbf{n} \otimes \mathbf{n})\mathbf{v}_i - \eta_{\perp}(\mathbf{v}_i \cdot \mathbf{n})\mathbf{n}, \tag{33}$$

where $\mathbf{n}$ is the normal direction to the wall of the local environment. Friction along the wall direction is controlled by the coefficient $\eta_{\|}$, while the normal friction is determined by $\eta_{\perp}$. Since we do not find direct evidence of PV-nucleus or PV-cortex coupling from experiments, we assume only particle-based repulsive forces between them.

#### 1.4 Time integration

The net force on particle $i$ consists of pairwise DPD forces from its neighboring particles $j \in \{r_{ij} < r_c\}$, forces from shape energy gradients, and forces from wall interactions. The movement of particle $i$ is governed by Newton's second law as

$$m_i \frac{\mathrm{d}\mathbf{v}_i}{\mathrm{d}t} = \sum_{j \in \{r_{ij} < r_c\}} \Big(\mathbf{F}_{C,ij} + \mathbf{F}_{D,ij} + \mathbf{F}_{R,ij}\Big) + \mathbf{F}_{\text{Shape},i} + \mathbf{F}_{\text{coupling},i} + \mathbf{F}_{\text{wall},i}. \tag{34}$$

For simulations based on Eq. (34), we use the Large-scale Atomic/Molecular Massively Parallel Simulator (LAMMPS), where a standard Velocity-Verlet time integration scheme is employed. The calculation of forces, i.e., Eq. (10), Eq. (32), and Eq. (33), is implemented in customized routines that are written in C++ and embedded in LAMMPS.

##### 1.5 Simulation units and scales

Throughout the simulations, we employ LJ reduced units, where the fundamental mass $m_0$, length $l_0$, energy $\epsilon_0$, and the Boltzmann constant $k_B$ are set to 1. By choosing representative physical values for these fundamentals, we can recover the real-world scale of any variable. In particular, the maximum cortical tension coefficient $\tau_\alpha$ has dimensions of energy per length,

$$[\tau_\alpha] = \frac{[\epsilon_0]}{[l_0]} = \text{force}.$$

Taking $\epsilon_0 \sim k_B T \approx 4.1\,\text{pN}\cdot\text{nm}$ and a typical DPD length scale $l_0 \sim 0.1\,\mu\text{m} = 100\,\text{nm}$, we obtain the characteristic surface-tension scale

$$\frac{\epsilon_0}{l_0^2} \sim \frac{4.1\,\text{pN}\cdot\text{nm}}{(100\,\text{nm})^2} \approx 0.41\ \text{pN}/\mu\text{m}.$$

Experimentally measured cortical tensions in amoeboid cells lie in the range 10 to 100 pN/$\mu$m [11, 12], so that our model's intrinsic tension scale is roughly 25–250 times smaller. We can place our simulations in the experimental regime by applying a scaling factor, e.g., 100, in the range. Under the mapping,

$$\frac{\epsilon_0}{l_0} \sim \frac{4.1\,\text{pN}\cdot\text{nm}}{100\,\text{nm}} \approx 0.041\ \text{pN},$$

so that $\tau_\alpha = 1$ in simulation units corresponds to $\approx 0.041 \times 100 = 4$ pN. Thus, by varying $\tau_\alpha \in [0, 2]$, we cover an effective force range of 0–8 pN, which is compatible with the lower end of experimentally reported cortical tensions once distributed around the cell periphery.

#### 2 Results

We focus on the question of whether the relative positions of the nucleus and the PV change under different conditions. Since we observed in experiments that the nucleus is more likely to be positioned in front of the PV before the cells entered the chambers, we set up our simulations with the same initial configuration. To quantify the relative position of the PV over time, we measure the fraction of reposition events in $N_{\text{repetition}}$ simulations as

$$Q = N_{\text{reposition}}/N_{\text{repetition}}. \tag{35}$$

Moreover, we quantify the relative position along the cell polarity axis with

$$\mathbf{r}_{\text{rel}} = \mathbf{r}_{\text{PV,front}} - \mathbf{r}_{\text{nucleus,front}}, \tag{36}$$

where $\mathbf{r}_{\text{PV,front}}$ and $\mathbf{r}_{\text{nucleus,front}}$ are defined based on the direction of motion $\hat{\mathbf{v}}_{\text{cell}}$ as

$$\mathbf{r}_{\text{PV,front}} = \arg\max_{\mathbf{r}_{\text{PV},i}}\big(\mathbf{r}_{\text{PV},i} \cdot \hat{\mathbf{v}}_{\text{cell}}\big), \tag{37}$$

$$\mathbf{r}_{\text{nucleus,front}} = \arg\max_{\mathbf{r}_{\text{nucleus,i}}}\big(\mathbf{r}_{\text{nucleus},i} \cdot \hat{\mathbf{v}}_{\text{cell}}\big). \tag{38}$$

To furthermore quantify the stress acting on each organelle, we adopt a coarse-grained Irving–Kirkwood approach and define a per-particle stress tensor as [13],

$$\mathbf{S}_i = -\frac{1}{V_i}\Big[m\,\mathbf{u}_i \otimes \mathbf{u}_i + \frac{1}{2}\sum_j \mathbf{r}_{ij} \otimes \mathbf{F}^C_{ij}\Big], \tag{39}$$

where $\mathbf{u}_i = \mathbf{v}_i - \bar{\mathbf{v}}_i$ is velocity of particle $i$ relative to the local mean fluid velocity $\bar{\mathbf{v}}_i$ computed within a circular environment determined by a cutoff $r_c$. The quantity $V_i = 1/\rho_i$ is the coarse-grained volume (area in 2-D) assigned to the particle $i$ via a density estimator

$\rho_i = \sum_j \frac{3}{\pi r_c^2} w(r_{ij})$. The conservative DPD force $\mathbf{F}_{ij}^C = a(1 - r_{ij}/r_c)\hat{\mathbf{r}}_{ij}$ enters the virial sum, while dissipative and random forces are excluded because they average to zero in the hydrostatic pressure and contribute only to the fluctuations. Finally, we extract a scalar pressure

$$p_i = -\frac{1}{2}\mathrm{tr}\,\mathbf{S}_i \tag{40}$$

for each particle, which can then be averaged over regions or used in strip-based force estimates on the nucleus and PV. To estimate the pushing force exerted by the fluid on each organelle, we define thin rectangular stripe regions near the front and back edges of each organelle, with perpendicular orientation to the cell polarity axis. We then calculate the average scalar pressure $\bar{p}$ within each stripe region and multiply it by the respective cross-sectional length $L_\perp$ of that region, which measures the organelle's transverse width. The net fluid force on an organelle is thus approximated as

$$\mathbf{F}_{\text{push}} = \left(\bar{p}_{\text{back}} L_{\perp,\text{back}} - \bar{p}_{\text{front}} L_{\perp,\text{front}}\right)\mathbf{n}, \tag{41}$$

where $\mathbf{n}$ is the unit vector along the cell polarity direction pointing from the back to the front of the cell. Positive values indicate a forward-directed push, while negative values imply a backward-directed pull.

#### 2.1 Model validation

To validate our model, we first compare the simulation results from cells containing only a nucleus with experimental observations in uninfected cells. As a measure for quantification of the nuclear position in a cell moving in the direction $\mathbf{n}$, we record the nuclear center-of-mass position $\mathbf{x}_{c,\text{nucleus}}$, the cell front $\mathbf{x}_{\text{front,cortex}}$, and the cell rear $\mathbf{x}_{\text{back,cortex}}$. Then, we define a normalized scalar as

$$x_{\text{nuc}} = \frac{\left(\mathbf{x}_{c,\text{nucleus}} - \mathbf{x}_{\text{back,cortex}}\right)\cdot\mathbf{n}}{\left(\mathbf{x}_{\text{front,cortex}} - \mathbf{x}_{\text{back,cortex}}\right)\cdot\mathbf{n}}. \tag{42}$$

where values close to unity indicate a front-positioned nucleus.

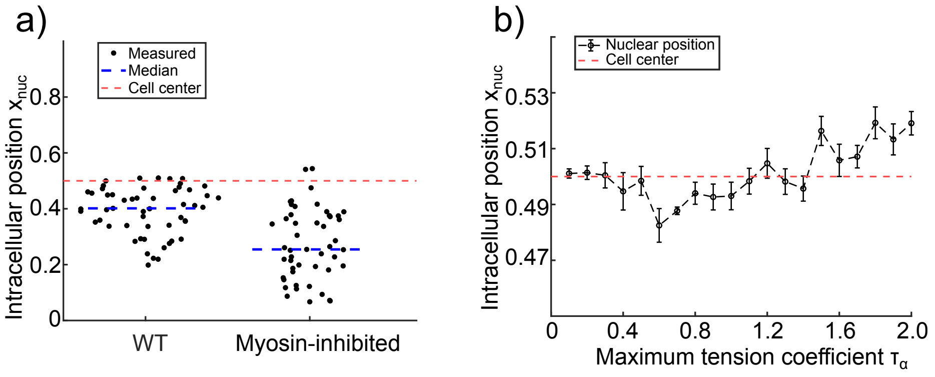


Figure 1: Comparison of nuclear positioning in experiment and simulation. (a) Distribution of the nuclear position measured in uninfected cells under control and myosin-inhibited conditions (see main text). (b) Average nucleus position in simulations with increasing cortical tension. Nuclear positions are measured at a fixed simulation time $t = 3 \times 10^4$. Error bars represent the standard error of the mean calculated from 10 separate simulation replicates for each value of $\tau_\alpha$.

Experimentally measured values of $x_{\text{nuc}}$, as shown in Fig. 1, for wild-type (left) and myosin-inhibited cells (right) show that the nucleus is closer to the cell rear in myosin-inhibited cells, suggesting that a myosin-dependent cortical contractility pushes the nucleus towards the cell front. Our simulations produce a similar behavior: at low cortical tension (weak contractility), the nucleus remains near its equilibrium position in the center of the cell. At intermediate tension, the nucleus shifts to the rear of the cell; at high cortical tension (strong contractility), the nucleus is pushed toward the cell front. We explain the central nuclear location at low tension by an insufficient cortical particle density gradient, leading to nearly random polymerization and depolymerization of the cortex and a weakly polarized cell. The weak cortical particle flow cannot advect intracellular matter persistently; thus, the nucleus remains bound near its equilibrium position by the nucleus-cortex center-of-mass interaction governed by Eq. (28). At intermediate tension, the pushing force of the internal fluid is weaker than the nucleus-cortex frictional coupling (Eq. (30)), resulting in a net backward movement of the nucleus. In contrast, high tension generates a strong hydrodynamic push that overcomes the backward friction, driving the nucleus toward the cell front. These simulation results qualitatively capture the essential features of nuclear positioning driven by myosin-dependent contractility observed in our experiments.

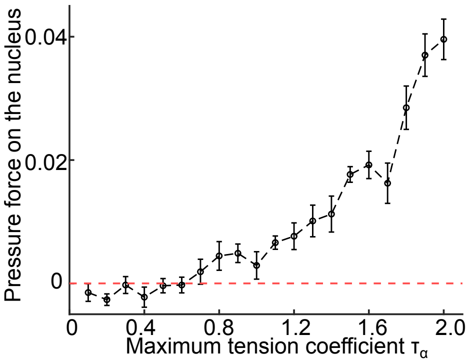


Figure 2: Mean pushing force on the nucleus $\langle F_{\text{nucleus,push}}(\tau_\alpha)\rangle_t$ averaged over all simulation time frames and all repetitions for each cortical tension value $\tau_\alpha$. Pushing forces are measured at simulation time $t = 3 \times 10^4$. Error bars represent the standard error of the mean calculated from 10 separate simulation replicates for each value of $\tau_\alpha$. A positive value of the pressure force, above the red line, indicates forward pushing.

Next, we quantify the pushing force on the nucleus by averaging across all simulation time frames for each cell. Figure 2 shows that the pushing force on the nucleus is close to zero at low cortical tension and increases with the cortical tension parameter $\tau_\alpha$, indicating that with stronger cortical tension, the nucleus experiences a greater pushing force from the intracellular medium, which shifts the equilibrium position of the nucleus toward the front of the cell. However, a coupling between the nucleus and the cortex can produce opposing forces, see Eqs. (28) and (30), which balance the pushing force from the internal fluid.

#### 2.2 Main results on PV positioning

We study how two key factors, cortical contractility and PV size, govern the repositioning. In our model, the myosin-driven contractility is represented by the cortical tension, Eq. (27), and governed by the parameter $\tau_\alpha$, and we control the PV size by varying its perimeter while keeping the nuclear perimeter constant. Figure 3a displays the reposition fraction $Q$ under various PV sizes and cortical-tension parameters $\tau_\alpha$. In general, repositioning is frequently observed in simulations of cells containing large PVs or with high cortical tension $\tau_\alpha$. Figure 3b shows the mean relative displacement $\langle x_{\text{rel}}\rangle = \langle x_{\text{PV,front}} - x_{\text{nuc,front}}\rangle$ in all repetitions with the current parameter set and reveals the same dependence on PV size and $\tau_\alpha$.
Experimental results show that larger PVs are correlated with increased myosin-driven cortical contractility. To investigate a corresponding interplay between PV size and cortical contractility in simulations, we sample along a straight line in this two-dimensional parameter space, as indicated by the white lines in Figs. 3a and b. We observe that as the PV

sizes increase, both the reposition fraction $Q$ and the mean relative distance $\langle x_{\mathrm{rel}} \rangle$ grow; see Fig. 3c. Then, we compute $\langle \Delta F(t) \rangle = \langle F_{\mathrm{PV,push}}(t) - F_{\mathrm{nucleus,push}}(t) \rangle$ at the sampling point $\star$ (Fig. 3a and b), where the ensemble average is computed among repositioned and unrepositioned simulations, respectively. Figure 3d shows that only cells with repositioned PVs exhibit $\langle \Delta F(t) \rangle > 0$ after repositioning is initiated. A similar separation is observed in all parameter sets of the phase plots (Fig. 3e), which confirms that a positive $\langle \Delta F \rangle$ is sufficient for the PV front movement. In summary, our model suggests that graded cortical tension generates a pressure-driven pushing force on the PV that, if exceeding the corresponding force on the nucleus, drives forward relocation. The PV size modulates this balance of mechanical forces by altering the cross-sectional area over which the pressure acts. The results align well with experiments along the diagonals of Fig. 3a and b, implying that myosin activity is upregulated by PV sizes *in vivo*.

#### 2.3 Role of model assumptions for PV positioning

##### 2.3.1 Nucleus-cortex coupling

To assess how each nucleus-cortex coupling contributes to PV repositioning, we perform three sets of simulations: (a) without the spring-like force positioning the nucleus at the center of mass , Eqs. (28) and (29), (b) without the frictional coupling, Eqs. (30) and (31), (c) with disabled coupling terms between the nucleus and the cortex. Figure 4 displays the fraction of simulations with repositioning and the mean relative displacement under the above conditions (a)-(c). A low frequency of repositioning events is observed for all sets of parameters. Additionally, the mean relative displacement remains negative regardless of the chosen parameters, confirming that the PV cannot overpass the nucleus under these conditions during the simulation time $t = 3 \times 10^4$. Thus, disabling either coupling term reduces the reposition fraction to near zero, confirming that both the center-of-mass tether and the frictional drag between the cortex and the nucleus are essential for the forward relocation of the PV. However, despite the fact that the disabling of the nucleus-cortex coupling prevents the repositioning of the PV in front of the nucleus, we continue to observe an influence of the size of the PV on its position. That is, for fixed $\tau_\alpha$, the PV moves further to the front of the cell as its size increases. This trend can be explained by the larger cross-sectional area onto which the fluid pressure generated in the rear of the cell acts.

##### 2.3.2 Mechanical properties

To examine how the chosen mechanical properties of PVs and nuclei in simulations affect their positioning inside the cell, we conduct the simulations with both the PV and the nucleus with the same number of particles and disable all the nucleus-cortex coupling. The simplest test is to only vary the perimeter elasticity constant $K_P$, while keeping other mechanical constants the same for PV and nucleus. In experiments, the nucleus appears softer and slimmer than the PV. To investigate an extreme case where the PV is ten times stiffer than the nucleus, we set $K_{P_{\mathrm{PV}}}/K_{P_{\mathrm{nuc}}} = 10$. The position of the PV is tracked in simulations of duration $t = 3 \times 10^4$ (default) and $t = 6 \times 10^4$ as shown in Fig. 5. The results show that, as is the case with the default parameters, repositioning is prevented in the default simulation time; Fig. 5a. However, repositioning can eventually occur after a sufficiently long time; see Fig. 5b. We conclude that a soft nucleus can facilitate PV positioning in the front to some extent.

##### 2.3.3 Influence of the cortex volume

Although line tension drives cortical flow, the resistance of the cortex to changes in area and perimeter, Eqs. (5) and (6), can affect how much intracellular pressure builds up. To study whether volume change alters model behavior, we remove the perimeter energy

penalty, Eq. (6), as an extreme case, allowing a more contractile cortex under the same line tension profile. We perform the same analysis as shown in Fig. 3. Figures 6a and b show that repositioning occurs more frequently with even lower values of PV sizes and cortical tension parameters $\tau_\alpha$, compared to Fig. 3. Additionally, increases in both measures, the reposition fraction $Q$ and the mean relative displacement $\langle x_{\text{rel}} \rangle$, along the diagonal parameter scan shown in Fig. 6a are steeper than in the default case, as shown in Fig. 6c. Compared to the default situation shown in Fig. 3e, removal of the volume constraint results in an earlier sign flip of the average pushing force difference $\langle \Delta F(t) \rangle$, indicating that the system starts to reposition the organelles earlier. This means that simulated cells without perimeter constraints are more prone to initiate repositioning by creating a pressure difference that flips the sign of $\Delta F(t)$. We explain this result with the observation that the cortex contracts more sharply under the same line tension without perimeter constraint, generating larger local pressure differences and thus triggering PV movement at lower tension or size thresholds. Nevertheless, the state diagram for PV repositioning remains qualitatively the same.
Together, these complementary results demonstrate that both the mechanical coupling and the mechanical properties of the organelles and the cortex influence the pressure-driven repositioning of the PV.

#### 3 Conclusions

We present a hybrid modeling framework that combines dissipative particle dynamics for intracellular hydrodynamics with a vertex-based representation for cell membranes and organelles. In the simulations, cell migration is driven by cortical flow, which is established by a graded cortical tension, as well as polymerization and depolymerization. We calculate the hydrostatic pressure using the Irving-Kirkwood stress tensor, allowing us to quantify the pushing forces on the two organelles.
We validated our model by simulating control cells with only the nucleus and no PV and reproduced the experimental observations of nuclear backward shifts when myosin is inhibited, confirming that a hydrostatic push is modulated by the graded cortical tension. Next, we showed that PV front repositioning occurs when the pushing force on the PV exceeds that on the nucleus. These forces are affected by cortical contractility, parameterized by $\tau_\alpha$, and the PV cross-section. The experimental results suggest that contractility increases with the PV size and, as a consequence, the larger PV is pushed in front of the nucleus. Finally, selected variations of the model assumptions demonstrated that, first, mechanical couplings between the nucleus and the cortex are crucial for PV repositioning, since they introduce a tendency to hold the nucleus at its equilibrium position. Second, constraints on organelle and cell sizes influence the results because a change in the size or the perimeter leads to a change in the pushing force on the PV, which in turn affects its repositioning. Together, these findings provide a minimal picture of the physical forces that determine the arrangement of the parasitic cargo relative to the nucleus in migrating immune cells.

[3] P. J. Hoogerbrugge and J. M. V. A. Koelman. Simulating microscopic hydrodynamic

phenomena with dissipative particle dynamics. *Europhysics Letters*, 19(3):155, jun 1992.

[11] Robert M Hochmuth. Micropipette aspiration of living cells. Technical report, 2000.

[12] R. Sakamoto and M. P. Murrell. Active tension and membrane friction mediate cortical flows and blebbing in a model actomyosin cortex. *Physical Review Research*, 6, 6 2024.

[13] John G. Kirkwood. The statistical mechanical theory of transport processes i. general theory. *The Journal of Chemical Physics*, 14(3):180–201, 03 1946.

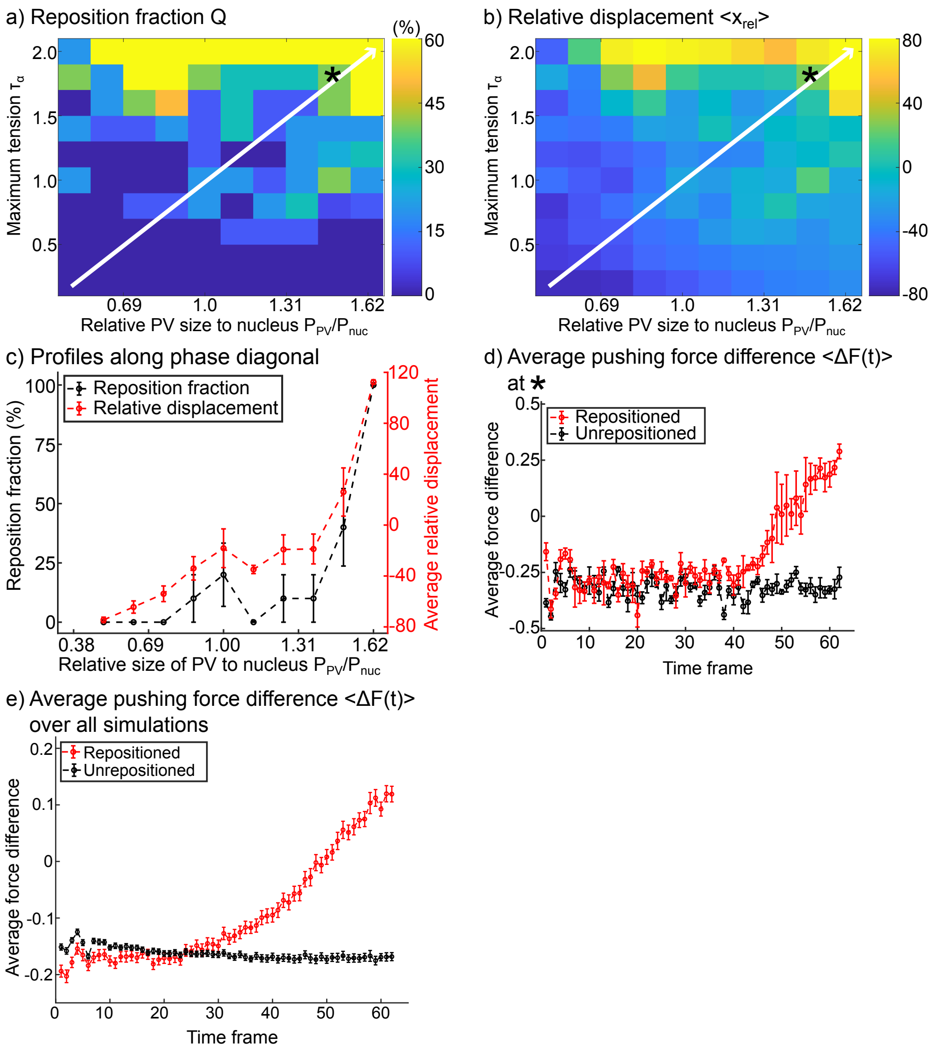


Figure 3: Repositioning of the PV under various simulation conditions. (a) Dependence of the reposition fraction $Q = N_{\text{reposition}}/N_{\text{repetition}}$ (in percent) on the cortical contractility parameterized by $\tau_\alpha$ and the PV size parameterized by the ratio of the PV's and nuclear perimeter $P_{\text{PV}}/P_{\text{nuc}}$. (b) Mean relative position $\langle x_{\text{rel}} \rangle = \langle x_{\text{PV,front}} - x_{\text{nucleus,front}} \rangle$. (c) Reposition fraction and mean relative position quantified along the straight white lines in parameter space shown in (a) and (b). (d) Average pushing force difference $\langle \Delta F(t) \rangle = \langle F_{\text{PV,push}}(t) - F_{\text{nucleus,push}}(t) \rangle$ between the PV and nucleus at every time frame in simulations at the sample point in parameter space labeled by $\star$ in (a) and (b). Ensemble averages are done by averaging over all time points in simulations where a PV repositioning either occurs or does not occur. (e) Average pushing force difference calculated from the same simulations as in (d). Here, ensemble averages are based on the instantaneous PV configuration. Timepoints in simulations are classified depending on whether or not the PV repositioning has occurred. Error bars represent the standard error of the mean in $\geq 10$ simulations.

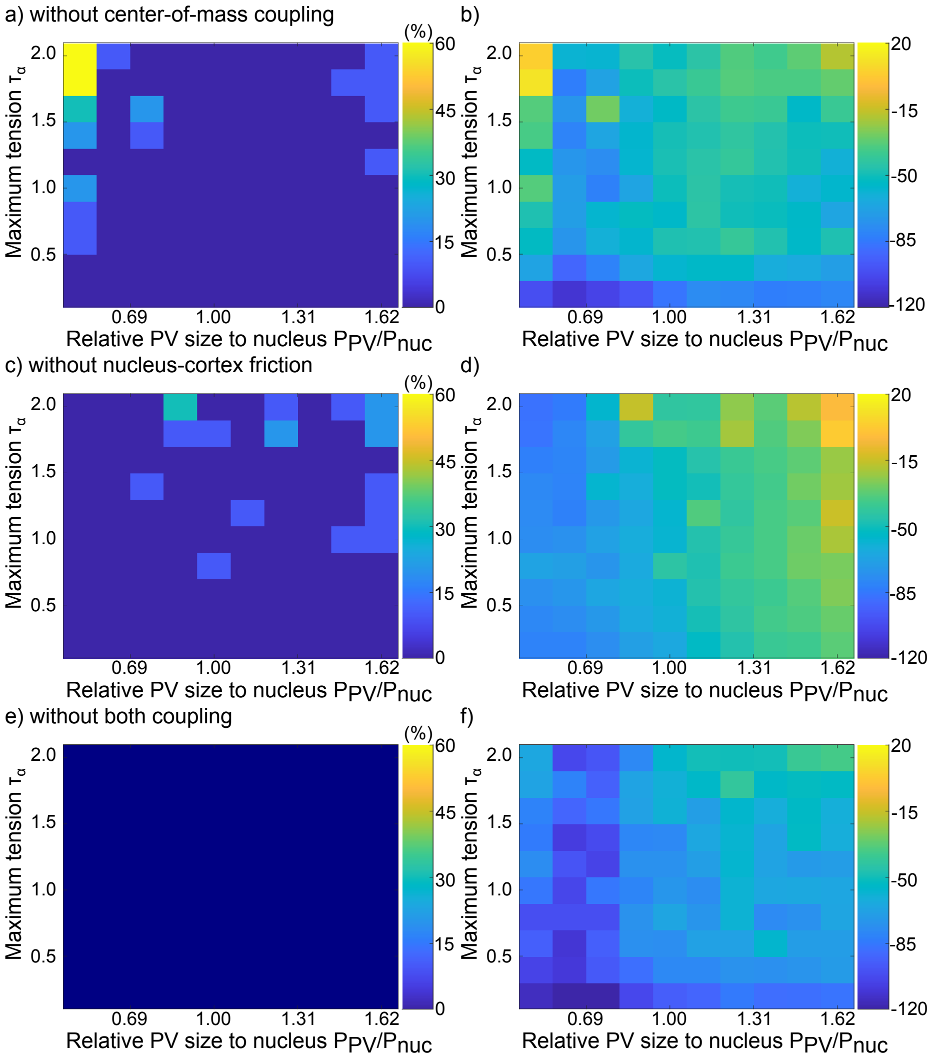


Figure 4: Reposition fraction $Q$ and mean relative displacement $\langle x_{\rm rel}\rangle$. (a)-(b) Only the nucleus-cortex center-of-mass spring is disabled. (c)-(d) Only the nucleus-cortex friction is disabled. (e)-(f) Both couplings are disabled. Turning off either interaction significantly suppresses the PV forward movement, suggesting the necessity of the nucleus-cortex mechanical coupling to observe reposition events within the simulation time $t = 3 \times 10^4$.

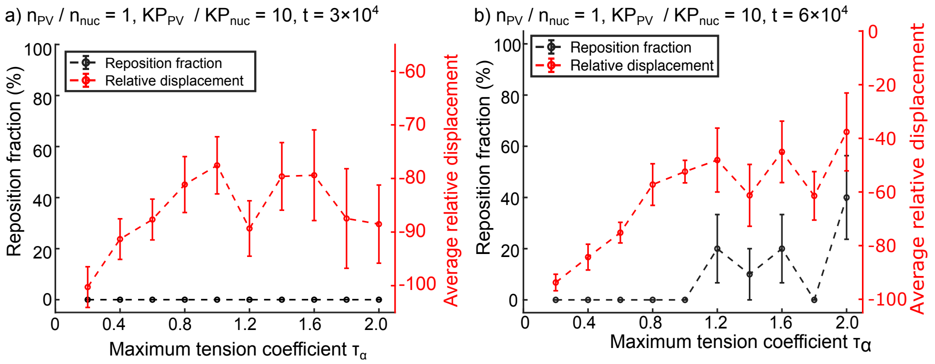


Figure 5: Reposition fraction $Q$ and mean relative displacement $\langle x_{\rm rel}\rangle$ for $K_{P_{\rm PV}}/K_{P_{\rm nuc}} = 10$. (a) Normal simulation time length. (b) Doubled simulation time. Error bars represent the standard error of the mean in $\geq 10$ simulations.

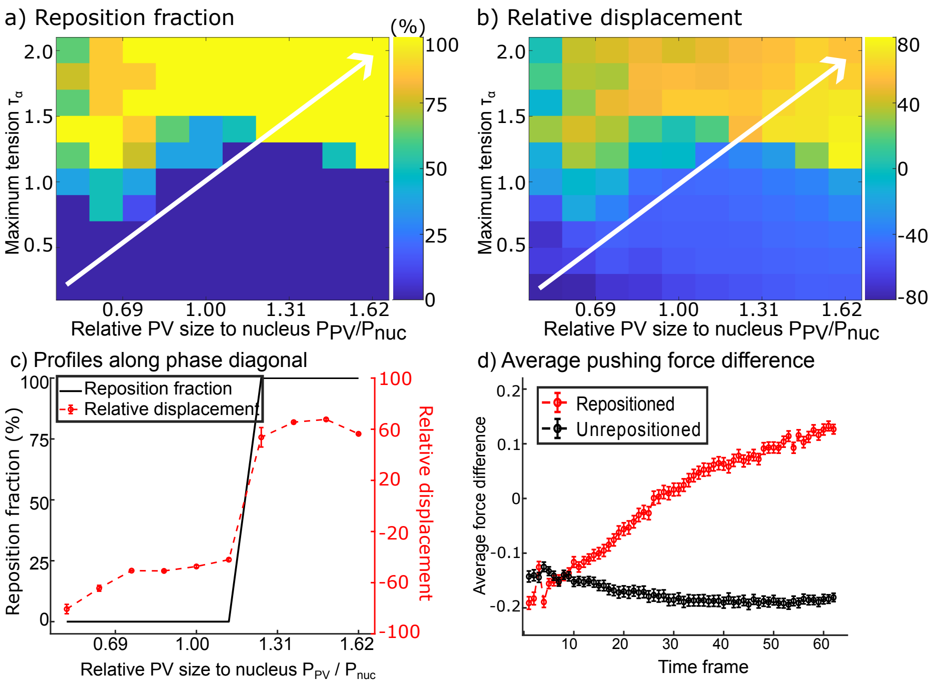


Figure 6: Results for simulations without a constraint on cortex perimeter change, facilitating a cell volume change with the tension parameter. (a) Reposition fraction $Q = N_{\text{reposition}}/N_{\text{repetition}}$ in percent. (b) Mean relative displacement $\langle x_{\text{rel}} \rangle = \langle x_{\text{PV,front}} - x_{\text{nuc,front}} \rangle$. (c) Repositioning quantified by $Q$ and $\langle x_{\text{rel}} \rangle$ along a path in parameter space indicated by the white lines in (a) and (b). (d) Average pushing force difference $\langle \Delta F(t) \rangle = \langle F_{\text{PV,push}} - F_{\text{nuc,push}} \rangle$ where the ensemble average is performed on repositioned and unrepositioned simulations. Error bars represent the standard error of the mean in $\geq 10$ simulations.
